# Far-red light absorption strategies and their structural basis in Photosystem I of Acaryochloris marina NIES-2412

**DOI:** 10.1101/2025.08.24.671928

**Authors:** Thomas J. Oliver, Eduard Elias, Giovanni Consoli, Ho Fong Leong, Violeta Cordón-Preciado, Andrea Fantuzzi, Tanai Cardona, A. William Rutherford, Roberta Croce

## Abstract

The marine cyanobacterium, *Acaryochloris marina*, uses the red-shifted chlorophyll *d* as its primary pigment, allowing it to absorb photons >700 nm. However, the widely studied type strain, *A. marina* MBIC11017, is atypical compared to most cyanobacteria, due to the absence of low energy chlorophylls (‘red forms’) within its Photosystem I complex. Consequently, Photosystem I and Photosystem II in the MBIC11017 strain share similar absorption spectra and are incapable of absorbing photons >740 nm. Recently, it has been discovered that the absorption and emission spectra from other *A. marina* strains are significantly more red-shifted than the MBIC11017 strain.

Here, we have combined advanced spectroscopy and high-resolution cryo-EM to characterize Photosystem I from *Acaryochloris marina* NIES-2412, a red-shifted strain that is more representative of the *A. marina* species. The structure resolves all 96 chlorophylls and cofactors and indicates the location of the red chlorophyll forms. Spectroscopic analysis reveals two distinct types of red forms: one arising from the classical mechanism of charge transfer–exciton mixing, and the other from purely excitonic interactions. Furthermore, we have identified PsaX2 as a critical subunit that fine-tunes the pigment geometries and energies to enable the formation of these red forms.

Together, these findings reveal how NIES-2412 PSI balances far-red light harvesting and energy trapping, highlighting its distinct strategy for adaptation in far-red light environments and redefining *A. marina* MBIC11017 as an atypical representative of the species.

## Introduction

While most cyanobacteria perform oxygenic photosynthesis using chlorophyll (Chl) *a*, *Acaryochloris marina* primarily uses the red-shifted Chl *d* to perform this process. Chl *d* differs from Chl *a* by the presence of a C3 formyl group rather than a vinyl group, resulting in a ∼30 nm red-shift. This enables *A. marina* to thrive in shaded environments, which are enriched in far-red light (FRL, 700 - 800 nm). While the enzyme responsible for the synthesis of Chl *d* is still unknown, the effect of its incorporation into the photosynthetic machinery has been the subject of a number of studies, primarily using the type strain *A. marina* MBIC11017 (Miyashita et al., 1996; Miyashita et al., 2003) (hereafter referred to as MBIC11017).

Photosystem I (PSI) is one of the multi-subunit pigment-protein complexes that make up the photosynthetic machinery of oxygenic phototrophs. It is formed by the two pseudo-symmetric core subunits, PsaA and PsaB, alongside 9 - 10 additional subunits, binding ∼95 Chls. The majority of these Chls act as antenna pigments, funnelling excitation energy to the reaction centre (RC), where charge separation leads to plastocyanin (or cytochrome c_6_) oxidation and ferredoxin reduction. In cyanobacteria, including MBIC11017, it is often present as a trimeric complex (Hamaguchi et al., 2021; Tomo et al., 2008; Xu et al., 2021).

MBIC11017 PSI exhibits both energetic and structural differences when compared to Chl *a*-containing PSI. The primary donor of MBIC11017 PSI is a Chl *d*/*d*’ dimer, P_740_, whose absorption is red-shifted by 40 nm compared to P_700_ in Chl *a* PSI. As such, there is ∼100 meV excited state energy loss associated with the incorporation of Chl *d* into MBIC11017 PSI. Interestingly, the P_740_ ^+^/P redox potential is essentially the same as the P_700_ ^+^/P_700_ couple (425 - 440 mV) (Bailleul et al., 2008; Schenderlein et al., 2008; Tomo *et al*., 2008), implying that the acceptor side cofactors of MBIC11017 PSI are energetically tuned to compensate for the P * free energy loss (Elias et al., 2024).

Structurally, the striking feature of MBIC11017 PSI is the presence of pheophytin (Pheo) *a* molecules in the A0 positions of the RC (Hamaguchi *et al*., 2021; Xu *et al*., 2021). Calculation of the free energy levels of the MBIC11017 PSI RC cofactors, based on ultrafast transient absorption measurements, estimate a ∼60 meV gap between P * and the P Pheo ^-^ radical pair, suggesting that Pheo is tuned to allow efficient charge separation (Petrova et al., 2023). However, Petrova *et al*. (2023) note that Chl *d* in the A0 position could also be tuned in a similar manner, hypothesising that the selective placement of Pheo *a* in this position is a structural requisite rather than an energetic one.

Despite these differences compared to Chl *a* PSI, there are surprisingly few consequences for the MBIC11017 PSI quantum efficiency of charge separation (i.e., its photon to electron conversion efficiency), which is ∼98% (Oliver et al., 2025), similar to Chl *a* containing PSI (Croce and van Amerongen, 2020). Time-resolved fluorescence measurements on MBIC11017 cells and isolated PSI complexes revealed a fast average energy trapping lifetime of 33.5 ps (Oliver *et al*., 2025). However, it was also observed that MBIC11017 does not contain any so-called red forms, i.e., Chl multimers absorbing at energies lower than the primary donor of the complex, in this case P_740_.

Red forms in PSI have been the subject of numerous studies (see Croce and van Amerongen (2013); Karapetyan et al. (2006) for reviews). Their characteristic far-red absorption occurs due to the strong excitonic interaction between two or more Chls, as well as contribution from charge-transfer (CT) states (Romero et al., 2009; Sláma et al., 2023; van Amerongen et al., 2000). The number, location and energies of these red forms varies depending on the organism. In cyanobacteria, Chl *f* containing PSI show red forms that emit at 800 nm (Nurnberg et al., 2018; Tros et al., 2021), Chl *a* containing PSI from *Arthrospira platensis* and *Nostoc punctiforme* possess red forms that emit up to 760 nm (Cardona and Magnuson, 2010; Karapetyan et al., 1997; Shubin et al., 1992) while *Gloeobacter violaceous* (Mangels et al., 2002) and *Synechococcus* sp. WH7803 (van Stokkum et al., 2013) PSI both lack red forms completely.

Due to their low energy, red forms act as local traps for excitation energy, necessitating uphill excitation energy transfer (EET) to the RC for charge separation to occur. The number and site energy of red forms within PSI roughly correlate with its overall trapping time (Gobets et al., 2001); in the absence of red forms, PSI trapping occurs in ∼14-18 ps (Gobets *et al*., 2001; van Stokkum *et al*., 2013), while the 760 nm emitting *A. platensis* PSI traps in ∼50 ps (Gobets *et al*., 2001; Karapetyan *et al*., 1997).

Although their influence on the energetic dynamics of PSI is clear, the function of red forms within PSI is less obvious. Several roles have been proposed, the simplest of which is to expand the PSI absorption spectrum into the far-red (Rivadossi et al., 1999; Trissl, 1993). This may be beneficial in low-light or shaded environments, or in dense culture environments where cell self-shading can be detrimental. Red forms have also been proposed as “concentrators” of excitation energy, acting as preferential pathways to the RC (Trissl, 1993).

The lack of red forms in MBIC11017 PSI is beneficial in terms of its quantum efficiency of charge-separation, but limits its capacity to absorb photons longer than 740 nm. This likely reflects MBIC11017 adaptation to a specific spectral niche, which is more enriched in white-light than far-red light (Oliver *et al*., 2025; Ulrich et al., 2021). Interestingly, it has recently emerged that there exists a large diversity of *A. marina* strains, many of which display red-shifted absorption and emission spectra *in-vivo* (Ulrich et al., 2024). Currently, the specific proteins and Chls responsible for the red-shifted absorption are unknown. However, PSI isolation from some of these strains has led to the suggestion that specific antenna proteins are responsible for their red-shift (Ulrich *et al*., 2024), in a manner analogous to algae and higher plants.

Here we have isolated the red-shifted Chl *d* containing PSI from one of these strains, *A. marina* NIES-2412 (hereafter referred to as NIES-2412). By combing a range of spectroscopic techniques (including ultra-fast time resolved fluorescence (TRF)), with cryo-electron microscopy (cryo-EM), we demonstrate that the NIES-2412 PSI complex is red-shifted due to a number of intrinsic red forms present in the core-antenna. We show that some of these red forms are homologous to the red forms found in many cyanobacteria, but one is novel and specific to certain strains of *A. marina*. The cryo-EM structure also revealed the presence of a highly divergent homolog of PsaX, that likely stabilises two of these red-shifted Chls. The presence of red forms in NIES-2412 PSI significantly extends its absorption into the far-red spectral region, up to ∼760 nm, but slows down its energy trapping.

## Results

### Spectroscopy of NIES-2412 PSI

Room temperature (RT) absorption spectra of NIES-2412 and MBIC11017 cells exhibit clear differences to one another (**Fig. 1A**). The NIES-2412 spectrum shows a shoulder to the main Q_Y_ absorption peak, extending its absorption to ∼760 nm. This shoulder is not present in MBIC11017. Additionally, the Q_Y_ absorption maximum of NIES-2412 (∼706 nm) is slightly blue-shifted compared to MBIC11017 (∼710 nm). Furthermore, NIES-2412 cells do not exhibit absorption in the 550-650 nm region, suggesting that they do not possess phycobilisome complexes, unlike MBIC11017.

**Figure 1.**
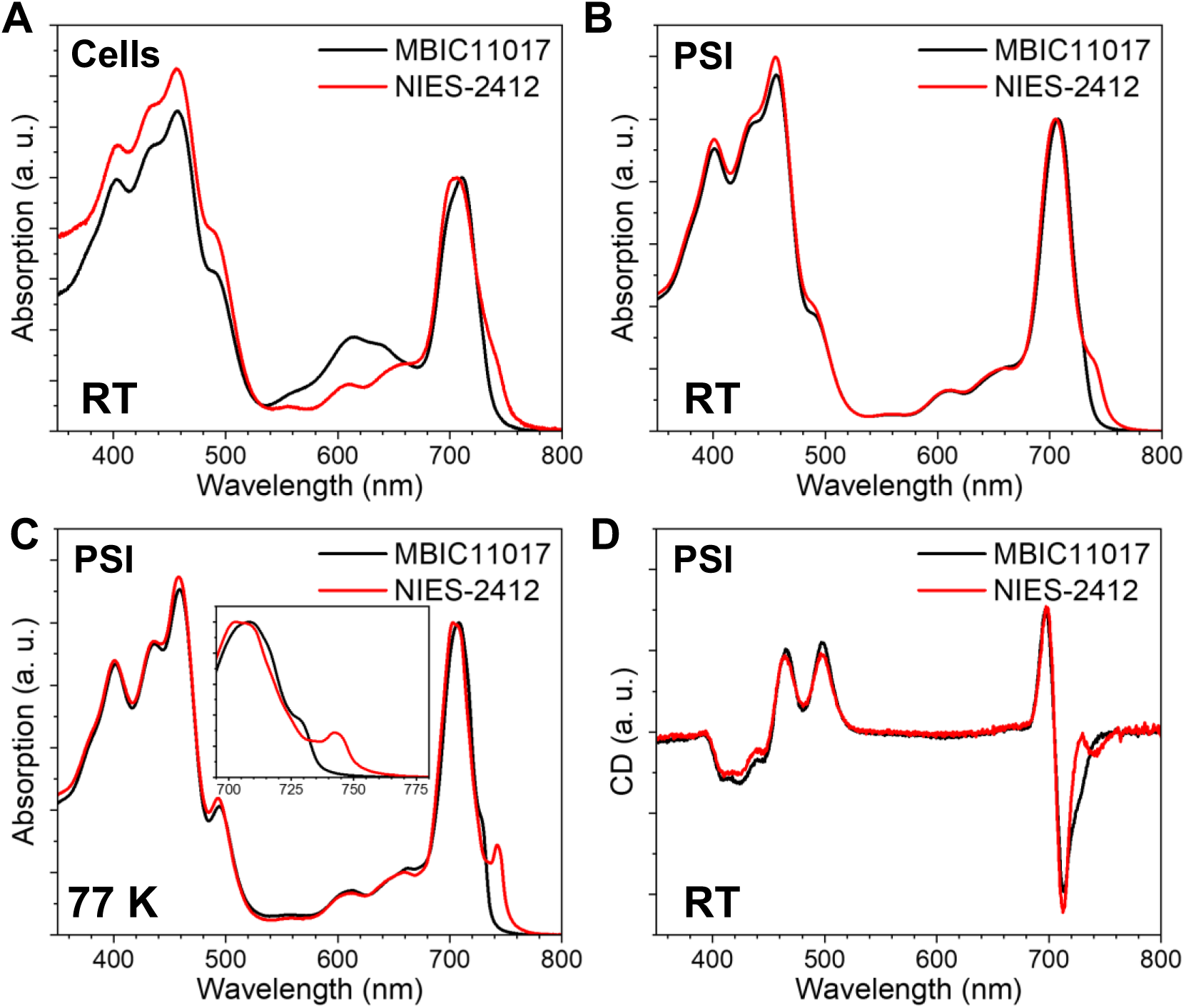
Absorption and circular dichroism spectra of *Acaryochloris marina* NIES-2412 cells and the isolated Photosystem I (PSI) complex compared to *Acaryochloris marina* MBIC11017. **(A)** Room temperature (RT) absorption spectra of NIES-2412 and MBIC11017 cells. Spectra were normalised to their maxima at ∼706-710 nm. **(B)** RT absorption spectra of NIES-2412 and MBIC11017 isolated PSI complexes. Spectra are normalised to their maxima at ∼708 nm. **(C)** 77 K absorption spectra of NIES-2412 and MBIC11017 isolated PSI complexes. Spectra are normalised to their maxima at ∼703-708 nm. Inset shows the Q_Y_ region of the spectrum between 705 and 775 nm. **(D)** Circular dichroism (CD) spectra of NIES-2412 and MBIC11017 isolated PSI complexes. Spectra are normalised to their integrated absorption spectrum between 670-800 nm.

To investigate the origin of the red-shifted shoulder, PSI from NIES-2412 was isolated (**Fig. S1**) and spectroscopically characterised. Like the *in-vivo* absorption spectrum, the RT absorption spectrum of NIES-2412 PSI (**Fig. 1B**) still exhibits the red shoulder. At 77 K, the red shoulder narrows (**Fig. 1C**), appearing as a distinct band with a maximum at 743 nm. Interestingly, the main Q_Y_ absorption peak of NIES-2412 PSI also differs from that of MBIC11017 PSI: it does not contain the small shoulder at 728 nm and its Q_Y_ maximum is blue-shifted relative to MBIC111017 PSI (∼703 nm vs ∼708 nm). The absorption spectra of pigments extracted from each PSI complex are essentially identical (**Fig. S2**), with maxima at 692.8 nm indicative of Chl *d*, although MBIC11017 PSI possess a small Chl *a* shoulder at 660 nm.

A gaussian deconvolution of the Q_Y_ region of the NIES-2412 PSI 77 K absorption spectrum was performed (**Fig. S3B**), guided by the 2^nd^ derivative of the spectrum (see **Methods and Fig. S3A)**. It shows that the red-shifted absorption peak can be predominately described by three Gaussians: one relatively narrow band centred at 744 nm (FWHM=∼8.5 nm), a broad one centred at 745 nm (FWHM=22 nm) and a third much broader one centred at 750 nm (FWHM=47 nm). Considering that the NIES-PSI complex contains 96 chlorins per monomer (see structural section, below), the areas under these Gaussians corresponds approximately to 3.6, 3.6 and 1.5 Chl *d* molecules. However, it is likely that the P_740_ absorption, which should correspond to an oscillator strength of ∼2 Chls *d* (Tomo *et al*., 2008), is also described by the 744 or 745 nm Gaussians.

Because the intrinsic circular dichroism (CD) signal of Chls is small, the CD spectrum in the visible region is dominated by excitonic interactions that occur between Chls (Garab and van Amerongen, 2009). The CD spectra of NIES-2412 PSI and the MBIC11017 PSI are shown in **Fig. 1D**. The spectra are very similar in the Soret region, but substantially different in the Q_Y_ region. The main difference is that the negative band at ∼710 nm in the NIES-2412 PSI spectrum lacks the shoulder that is present in the MBIC11017 PSI spectrum and instead possesses a distinct negative peak at ∼742 nm, which is not present in MBIC11017. This latter feature is clearly indicative of an excitonic interaction that is present in NIES-2412 PSI and absent in MBIC11017 PSI, with the lower energy band absorbing around 740 nm.

The RT emission spectrum of NIES-2412 PSI is red-shifted relative to MBIC11017 PSI (**Fig. 2A**) and consists of three components: a shoulder at ∼720 nm (the maximum of the MBIC11017 emission), a peak at 744 nm, and a broad red tail that extends past 800 nm. At 77K, the NIES-2412 PSI emission displays a maximum at 767 nm, which is red-shifted by 36 nm compared to the MBIC11017 maximum (**Fig. 2B**). Additional bands at ∼700 nm and 750 nm are also present in the 77 K emission spectrum, but these are likely to originate from a small contamination of free Chl *d* and disconnected antennae which have much longer fluorescence lifetimes than the NIES-2412 PSI (see time-resolved fluorescence results, below), and hence are disproportionately represented in the steady-state emission spectrum.

**Figure 2.**
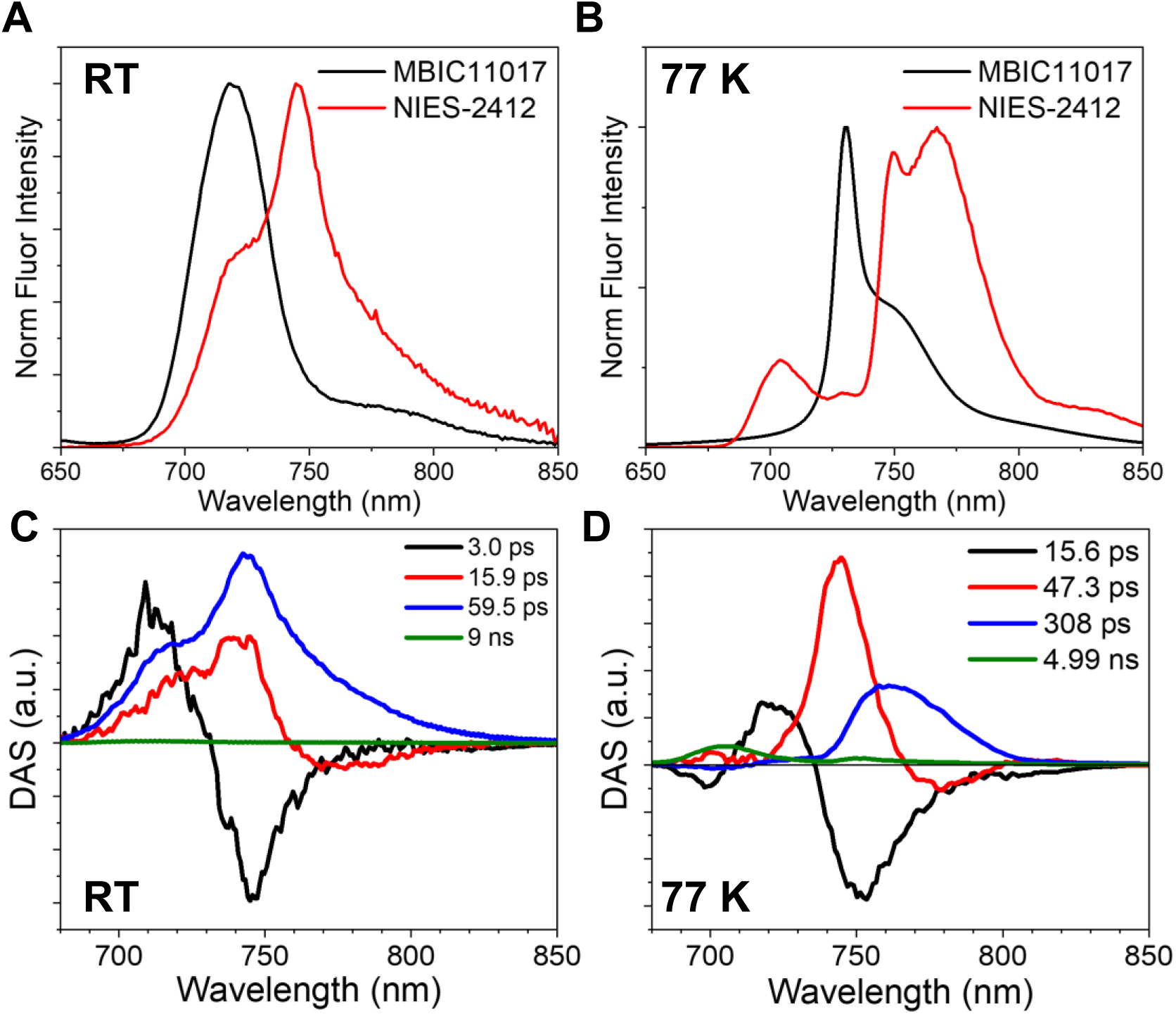
Steady-state and time-resolved fluorescence (TRF) spectra from *Acaryochloris marina* NIES-2412 Photosystem I (PSI). **(A)** Room temperature (RT) fluorescence emission spectra of NIES-2412 and MBIC11017 (taken from Oliver *et al*. (2025)) isolated PSI complexes. Spectra were normalised to their emission maxima. **(B)** 77 K fluorescence emission spectra of NIES-2412 and MBIC11017 (taken from Oliver *et al*. (2025)) isolated PSI complexes. Spectra were normalised to their emission maxima. **(C)** Global analysis of RT time-resolved fluorescence data from the NIES-2412 isolated PSI complex. **(D)** Global analysis of 77 K time-resolved fluorescence data from the NIES-2412 isolated PSI complex.

To investigate the effect of the red-shifted chlorophylls in NIES-2412 PSI on its excitation-energy trapping dynamics, we measured its fluorescence decay using a streak camera setup with 400 nm excitation. Decay associated spectra (DAS) were obtained from global analysis of the RT TRF data (**Fig. 2C**). The RT dynamics show two components that possess both positive and negative amplitudes, indicative of EET processes. The first component has a lifetime of 3 ps and describes EET to a pigment pool emitting at ∼745 nm. The second component is longer (∼16 ps) and describes EET to a pigment pool emitting at ∼770 nm. This component is largely non-conservative, possessing a positive integrated area, indicating that energy trapping also occurs with a similar time constant. The main energy trapping component occurs in ∼60 ps, almost twice as long as observed in MBIC11017 PSI (Oliver *et al*., 2025). The spectrum of this component is very broad, possessing the same shape as the RT emission spectrum. A very small, nanosecond component was also resolved, arising from disconnected Chl *d*.

At 77 K, the global analysis of TRF data also shows four components (**Fig. 2D**). The first two represent EET to Chl *d* pools emitting at ∼750 nm and ∼770 nm and have lifetimes of ∼15 ps and ∼50 ps, respectively. As was the case at RT, the second component is largely positive, indicating that excitation energy trapping from the 750 bands is also occurring within 50 ps. The third component, with a lifetime of ∼300 ps, is positive, very broad and peaks at ∼765 nm, indicating trapping from a very low energy form. The final component has a ∼5 ns lifetime and is very small in amplitude. It can be attributed to disconnected Chls d and antenna proteins.

The excitation energy dynamics of numerous Photosystem I complexes can be described by a kinetic scheme in which excitation energy is trapped from a large number of bulk Chls, which are in thermal equilibrium with one or more red forms (Croce and van Amerongen, 2013; Gobets and Van Grondelle, 2001; Gobets *et al*., 2001). To assess the validity of such a model, a Stepanov analysis can be performed (Jennings et al., 2003; Stepanov, 1957): if there is a good match between the measured emission spectrum and the emission spectrum as determined from the application of the Stepanov relation to the corresponding absorption spectrum, then this is a clear indication of the excitation energy being trapped from a thermally equilibrated state. The measured emission spectrum of NIES-2412 PSI and its predicted spectrum based on the Stepanov relation are shown in **Fig. S4**. A close match between the two spectra is observed, the main difference being ∼720 nm, which can be explained by some impurity of the sample. The clear similarity between the predicted and measured emission spectrum demonstrates that a target kinetic scheme in which bulk Chls are connected to the two separate red forms is valid (Jennings *et al*., 2003).

The kinetic scheme and fitted parameters of the target analysis are shown in **Fig. 3A**, and its associated SAS in **Fig. 3B**. Two red forms are clearly disentangled from the data, the first one peaking at 745 nm (Red 1) and the second one at 757 nm (Red2). The Bulk SAS is relatively broad and is very similar to the RT emission spectrum of MBIC1107 PSI (**Fig. S5**). Trapping from these Bulk Chls occurs with a characteristic time constant of 20.4 ps in NIES-2412 PSI.

**Figure 3.**
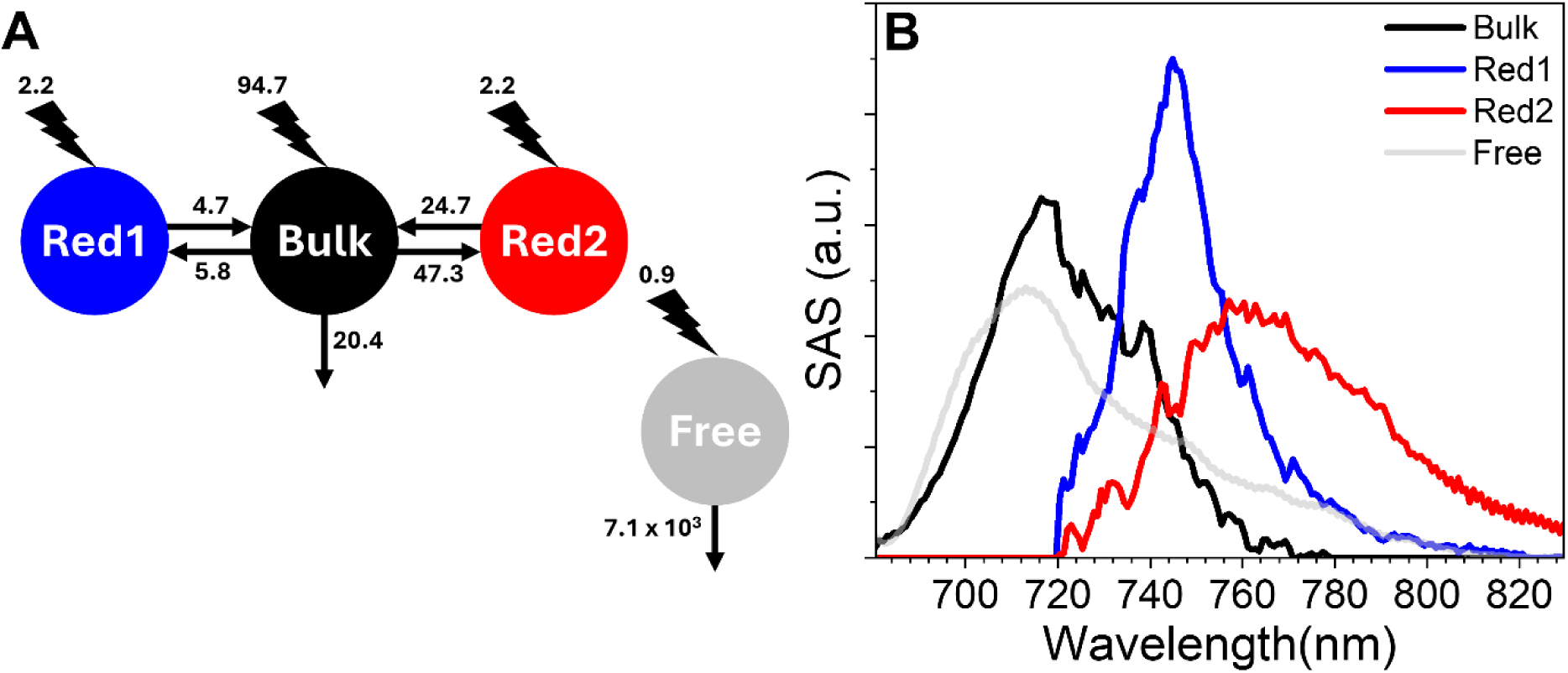
Target analysis results for the NIES-2412 PSI RT time-resolved fluorescence data. **(A)** Target kinetic scheme. The numbers above the lightning bolts indicate the fitted portion of initial excitation for the different compartments. The numbers connected to the arrows indicate the reciprocal of the fitted kinetic rates in ps. **(B)** SAS for the fitted kinetic scheme. The Free compartment SAS has been smoothed for clarity.

As observed in the global analysis (**Fig. 2C**), the Red1 compartment is populated quickly from the Bulk compartment (5.8 ps), whereas the Red2 compartment is populated on a slower timescale (47.3 ps). Due to the far fewer Chls contributing to the Red1 and Red2 compartments than to the Bulk compartment, the equilibrium between the Bulk and red forms is shifted towards the Bulk Chl pool, despite its higher energy. Considering these equilibria, the peak positions of the SAS, and the total number of Chls per PSI, the number of Chls in each compartment are estimated to be ∼5 Chls for Red1 and ∼1 for Red2 (see **Methods** and **Tab. S1**). These values are very close to the estimates from the Gaussian deconvolution (6.7 Chls *d* absorbing above 740 nm, not including P_740_).

### Structure of NIES-2412 PSI

To understand the molecular origin of the Red1 and Red2 pools, the NIES-2412 PSI structure was solved through Cryo EM single particle analysis, yielding a map of trimeric PSI (refined with C3 symmetry) at a Gold Standard Fourier Shell Correlation (GS-FSC) resolution of 2.64 Å (**Fig. S6, Tab. S2**).

The map reveals electrostatic potential for 12 subunits coordinating a number of cofactors: 93 Chls *d*, 1 Chl *d*’, 21 α-carotenes, 2 pheophytins *a*, 2 phylloquinones, 3 Fe_4_S_4_ iron-sulfur clusters, as well as various lipids and water molecules. The total number of chlorins (96), is higher than those resolved in the two available structures of MBIC11017 PSI (73 (Hamaguchi *et al*., 2021) and 79 (Xu *et al*., 2021)), but very close to the MBIC11017 PSI biochemical estimate (100 (Tomo *et al*., 2008)). The number of chlorins resolved is also the same as in the *Thermosynechococcus elongatus* BP-1 (*T. elongatus*) PSI crystal structure (Jordan et al., 2001), although NIES-2412 PSI does not possess Chls *d* at similar positions to Chl *a* J1303 (ligated by H39 at the C-terminus of PsaJ) and Chl *a* M1601 (found at the monomer-monomer interface near PsaM). Instead, NIES-2412 PSI binds an additional Chl *d* in PsaB via D310 and another Chl *d* at the monomer-monomer interface next to PsaI.

Compared to the highest resolution MBIC11017 PSI structure (Hamaguchi *et al*., 2021), NIES-2412 PSI binds an additional 23 Chls *d*, 14 of which are bound by PsaB (**Fig. 4A**). In both reported MBIC11017 PSI structures, the peripheral loops of PsaB, between the 4^th^ and 5^th^ helices on the cytosolic side of the membrane, and the 7^th^ and 8^th^ helices on the luminal side, are un-resolved. The authors attribute this to flexibility within these regions due to the absence of the PsaX subunit in MBIC11017 PSI. The structure of NIES-2412 PSI reveals a single transmembrane helix subunit that is not present in either MBIC11017 PSI structure. To obtain an improved ESP for this subunit, symmetry expansion of the PSI trimer followed by local refinement was performed, resulting in a map at a resolution of 2.44 Å. This subunit interacts with both the cytosolic and luminal PsaB loops, likely stabilising them and facilitating the resolution of both the loops and their associated Chls *d*. Although this subunit appears very different to PsaX from other cyanobacteria (20 % sequence identity vs. *T. elongatus* PsaX), its similar position within the PSI structure (**Fig. 4B**) supports its identification as PsaX. We therefore refer to it as PsaX2 throughout the manuscript.

**Figure 4.**
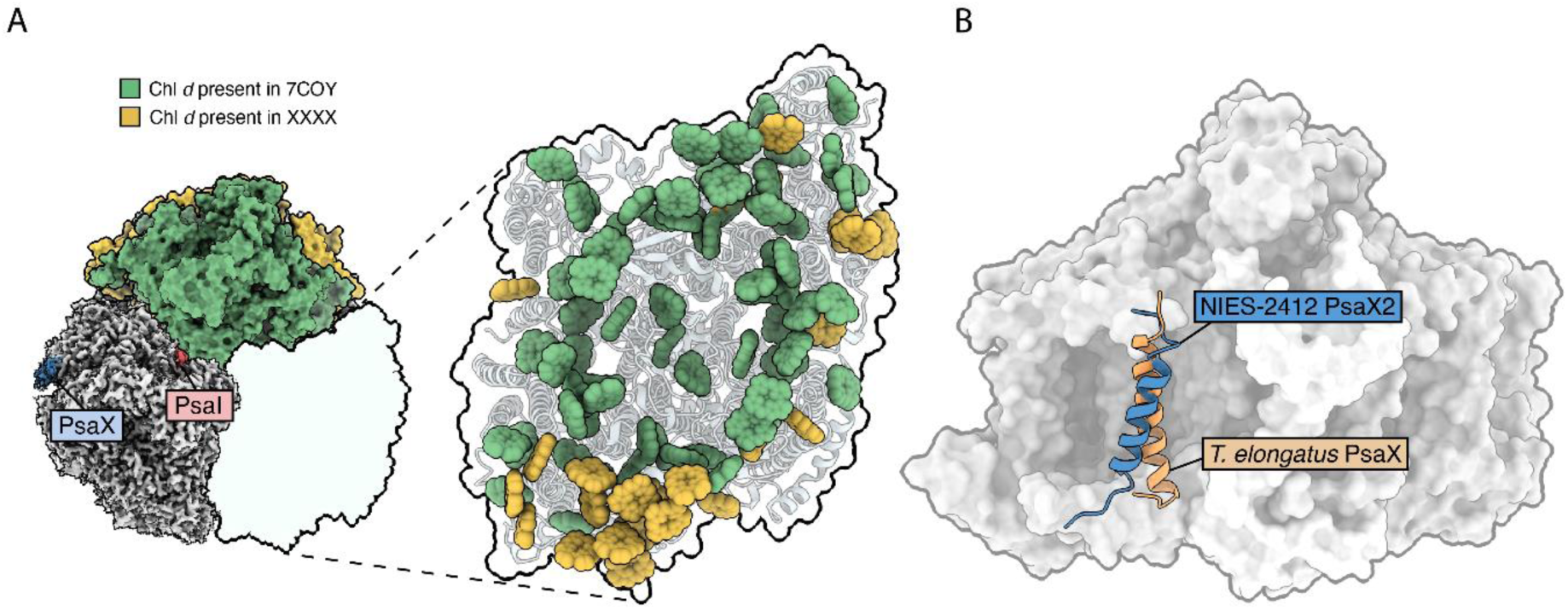
The structure of the trimeric Photosystem I (PSI) complex from *Acaryochloris marina* NIES-2412, showing the location of all resolved Chl *d*, in comparison with *Acaryochloris marina* MBIC11017 PSI. **(A)** View of the trimeric NIES-2412 electrostatic potential (ESP) map from the cytoplasmic side. The map is shaded in yellow to show the additional areas that were resolved in the NIES-2412 PSI monomer, relative to MBIC11017 PSI (PDB ID: 7COY). Green shading shows common areas, which were resolved in both structures. The ESP map of the NIES-2412 PSI monomer is also shown, with PsaX and PsaI labelled. The zoomed in monomer shows the locations of the Chls within the NIES-2412 PSI monomer, with those shaded in green present in both the NIES-2412 and MBIC11017 structures and those shaded in yellow only present in NIES-2412. **(B)** View of the ESP map from the plane of the membrane, looking at PsaB, showing the position of PsaX2. PsaX2 is coloured blue, with *T. elongatus* PsaX overlaid in orange to show their relative positions.

#### Candidate Red Forms In NIES-2412 PSI

Given that red forms originate from electronic coupling between two or more Chls (Gobets and Van Grondelle, 2001; Morosinotto et al., 2003; Romero *et al*., 2009) we initially searched for Chl *d* dimers or trimers within the additionally resolved ESP within PsaB. This yielded two contenders for the NIES-2412 PSI red forms (**Fig. 5A**), both of which interact with PsaX2. The first is the B31-B32-B33 Chl *d* trimer, bound by the loop between the 7^th^ and 8^th^ helices (**Fig 5E**). The first Chl within this trimer (B31) is co-ordinated by PsaB His^465^, whereas the second (B32) and third (B33) are both coordinated by water molecules, the latter of which is H-bonded by PsaB Asn^489^. Additionally, PsaX2 Trp^26^ also provides a H-bond to the C3 formyl group of Chl *d* B33. The separation between these Chls *d* is small, with centre-to-centre distances of ∼8.4 Å, and their Q_Y_ transition dipole moments are essentially parallel.

**Figure 5.**
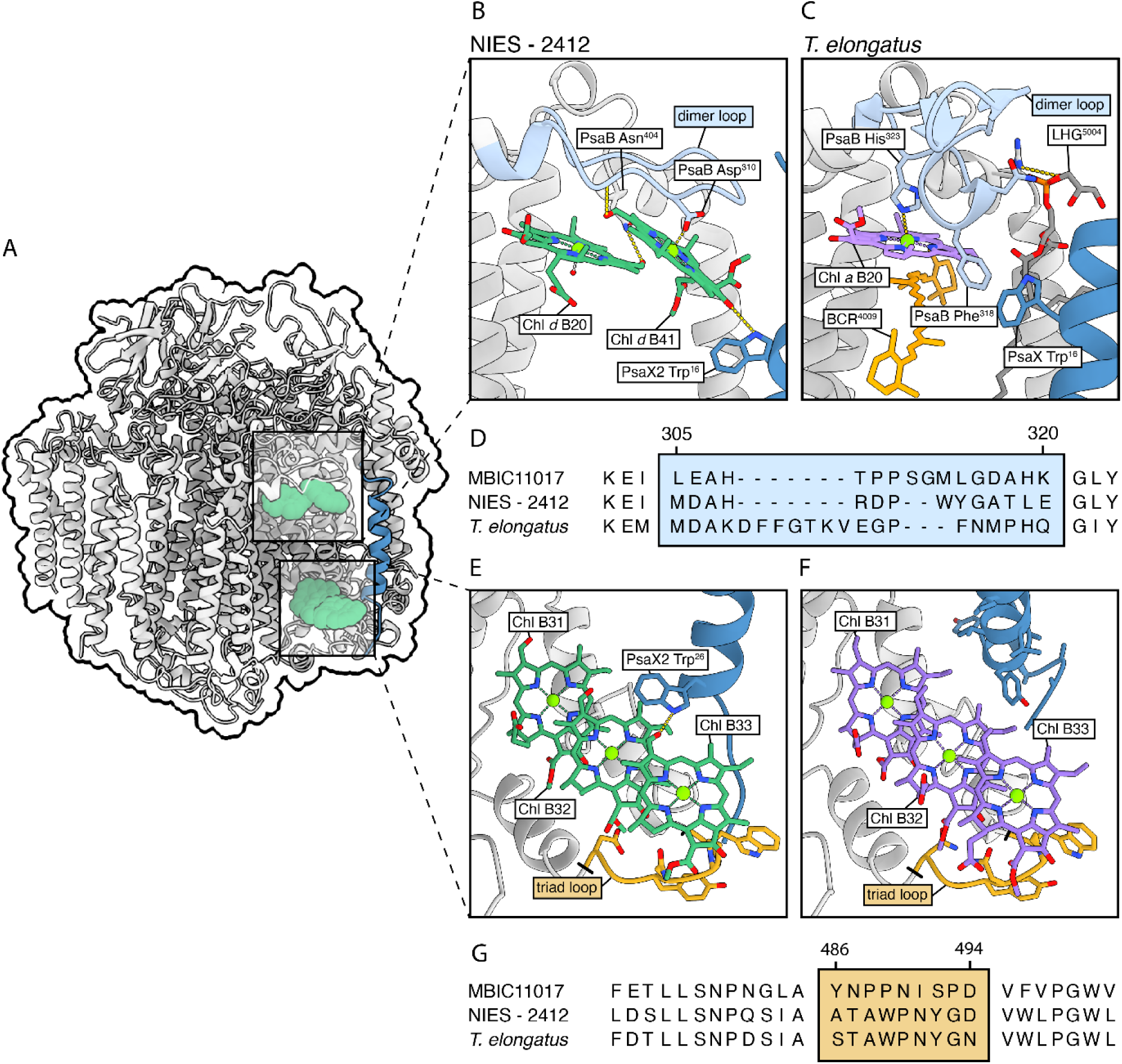
The location of the proposed additional red forms in Photosystem I (PSI) from *Acaryochloris marina* NIES-2412. **(A)** View of the NIES-2412 PSI monomer parallel to the thylakoid membrane showing the location of the two proposed red forms on the cytosolic and luminal sides of PsaB (shaded in green), and the single transmembrane helix, PsaX2, that interacts with them both, shaded in blue. **(B)** The B20-B41 Chl *d* dimer bound by PsaB in NIES-2412. The B41 Chl *d* is ligated by PsaB Asp^310^ (pale blue loop), and receives a H-bond from PsaX2 Trp^16^ (to its C13^1^ carbonyl oxygen). The B20 Chl *d* is ligated by a water molecule and is H-bonded by PsaB Asn^404^ (to its C3 formyl group). **(C)** The same region within *T. elongatus*, showing the absence of a Chl *a* dimer. **(D)** Sequence alignment of PsaB in this region. **(E)** The B31-B32-B33 Chl *d* trimer bound by PsaB in NIES-2412. The B33 Chl *d* is ligated by a water molecule, which is H-bonded by PsaB Asn^489^ (orange loop). The C3 formyl group of the B33 Chl *d* is H-bonded by PsaX2 W26. **(F)** The B31-B32-B33 Chl *a* trimer in *T. elongatus*, also showing the PsaB loop and PsaX. **(G)** Sequence alignment of PsaB in this region.

The second candidate red form is the B20-B41 Chl *d* dimer, bound by the loop between the 4^th^ and 5^th^ helices of PsaB, on its cytosolic side (**Fig. 5E)**. The B41 Chl *d* is ligated by PsaB Asp^310^, and its C13^1^ carbonyl oxygen is hydrogen bonded by PsaX2 Trp^16^. Its partner, the B20 Chl *d*, differs from Chls found at this position in other cyanobacterial PSI structures due to its lack of ligation by a histidine residue. Instead, it is coordinated by a water molecule, and a leucine is present at the position normally occupied by PsaB His^322^. The B20 Chl *d* is H-bonded by Asn^404^ to its C3 formyl group. Like the B31-B32-B33 Chl *d* trimer, the B20-B41 dimer Chls *d* are positioned close to one another, with an inter-pigment separation of ∼8.4 Å and an almost parallel orientation of their Q_Y_ transition dipole moments.

To assess the degree of exciton delocalization within both the B31-B32-B33 trimer and the B20-B41 dimer, their electronic coupling values were calculated and exciton Hamiltonians diagonalized, assuming equal site energies (**Tab. S3-6**). This reveals that the lowest energy exciton of the trimer and the dimer have energies of 151 cm^-1^ and 122 cm^-1^ below that of their individual pigments, respectively. Furthermore, their dipole strengths are almost entirely distributed to these levels (98% and 93%, respectively).

Sequence alignment of PsaB from NIES-2412 and MBIC11017 reveals key differences in the B31-B32-B33 (**Fig. 5G**) and the B20-B41 (**Fig. 5D**) binding regions. In the case of the trimer, the MBIC11017 PsaB sequence is highly divergent, lacking the residue that H-bonds the water molecule that coordinates the third, outmost, Chl *d*. A similar scenario is observed for the B20-B41 dimer: the MBIC11017 sequence is highly dissimilar from the NIES-2412 sequence as it does not possess the B41 co-ordinating residue, Asp^310^, and the B20 Chl is likely co-ordinated by His^317^ (Leu in NIES-2412), instead of a water molecule, based on a histidine residue in this position in other cyanobacterial PSI structures (Jordan *et al*., 2001; Malavath et al., 2018). To summarise, we assign the B31-B32-B33 and the B20-B41 clusters to red forms.

However, their combined oscillator strength is ∼5. Considering that the Gaussian deconvolution of the 77 K absorption spectra suggests that ∼7 Chls *d* are responsible for the absorption above 740 nm, (not including P_740_), ∼2 more red Chls need to be assigned in NIES-2412. One previously suggested red form is the A12-A14 *(Jiang et al., 2025; Kato et al., 2022).* In NIES-2412 PSI, these Chls *d* are at a centre-centre distance of ∼8.4 Å and they are co-ordinated by PsaA His216 and a water molecule that is H-bonded by PsaA His241. While both residues are present in PsaA from MBIC11017, a number of residues surrounding the NIES-2412 A12-A14 dimer are absent in MBIC11017 (**Fig. S7**). These residues may stabilise the A12-A14 dimer in NIES-2412, allowing for a stronger electronic interaction between the two Chls *d*. Indeed, the A12-A14 dimer is not resolved in the MBIC11017 structure published by Hamaguchi *et al*. (2021) (although it is present in the structure of Xu *et al*. (2021)).

Additional red forms candidates have previously been suggested in cyanobacterial PSI, including the B37-B38 and the B07-A32 dimers (Byrdin et al., 2002; Jordan *et al*., 2001; Khmelnitskiy et al., 2020; Riley et al., 2007). Comparison between the NIES-2412 and MBIC11017 structures shows that both organisms possess Chls *d* in these positions, with no differences in their ligands. However, in the case of the NIES-2412 B37-B38 dimer, there are amino acid changes around these Chls in PsaA, PsaB and PsaL (relative to MBIC11017), that could lead to altered electronic interactions between the two Chls *d* (**Fig. S8**). A similar scenario is found in the B07-A32 dimer, whereby the Chl co-ordinating residues are the same in both organisms, but there are multiple nearby residue changes in NIES-2412 PSI compared to MBIC11017.

#### An Acaryochloris specific PsaI-bound Chl *d*

The NIES-2412 PSI structure shows the presence of an additional Chl *d* (I01), coordinated by PsaI and located between PsaB of one monomer and PsaL of the adjacent one (**Fig. S9**). There is no Chl present at this position in any resolved Chl *a*-PSI structure. Inspection of the *T. elongatus* PsaI and PsaL sequences reveals that Chl binding at this position is blocked primarily by the presence of PsaI Trp^20^, as well as PsaL Met^149^. In NIES-2412 PsaI, there is an alanine instead of a tryptophan at this position, and the PsaL C-terminus is truncated, allowing the binding of Chl *d* here. These changes are also present in MBIC11017 (**Fig. S10**), and indeed a Chl *d* is observed in the same position in the structure of Xu *et al*. (2021).

Given the close proximity of this Chl *d* to the monomer-monomer interface, and the conservation of this site across different *Acaryochloris* strains, it could be argued that this Chl may play a role in inter-monomer EET in NIES-2412 PSI. To test this hypothesis, we calculated the excitation energy transfer rate between this Chl and its nearest neighbour in an adjoining monomer (i.e. Chl L03) using the framework laid out in Croce and van Amerongen (2020) (taking the parameters for Chl *a*→Chl *a* EET). The calculated energy transfer rate between these Chls is very small (0.02 ps^-1^), which would not allow for efficient EET across monomers, given that this Chl is much more strongly connected to other Chls within its own monomer (the calculated energy transfer rate from Chl I01 to Chl B07 is 2.9 ps^-1^). Interestingly, the centre-to-centre distance between the I01 and B07 Chls *d* is only 11.7 Å. It is possible that the presence of I01 could alter the electronic interaction between the B07-A32 Chls *d* (a potential red form).

## Discussion

### PSI in NIES-2412 is ∼20 nm more red-shifted than in MBIC11017

*Acaryochloris* is the only known genus of cyanobacteria that solely uses Chl *d* to absorb far-red light. In the most studied strain, MBIC11017, PSI contains a minimal number of red forms, and its ability to absorb FRL is mainly due to the intrinsically red-shifted nature of Chl *d*. The NIES-2412 PSI RT absorption spectrum extends to the far-red by ∼20 nm compared to MBIC11017, up to ∼760 nm, allowing it to absorb significantly more photons in the far-red region (700 - 800 nm).

Low temperature absorption spectroscopy shows that this arises due to 3 additional absorption bands in the 740 - 760 nm region, originating from ∼9 Chls in total, although the P_740_ Chl *d*/Chl *d*’ pair is likely responsible for 2 of these. Solvent extraction of the NIES-2412 and MBIC11017 PSI pigments did not reveal the presence of additionally Chls that are intrinsically red-shifted (such as Chl *f* (Chen et al., 2010)), suggesting that pigment-pigment and pigment-protein interactions involving Chl *d*, which are not present in MBIC11017, are responsible for this extra absorption capability above 740 nm.

The presence of chlorophylls absorbing at lower energy than the reaction centre (red forms) is a hallmark of most Chl *a*-containing PSI complexes and allows them to use far-red photons to some extent. In plant and algal PSI, the major red forms are typically located within antenna proteins, whereas in cyanobacteria they are integrated into the core itself. However, some cyanobacteria also utilise PSI antenna proteins, such as IsiA (Bibby et al., 2001; Boekema et al., 2001) or Pcb proteins (Bibby et al., 2003). The structure of NIES-2412 PSI reveals a trimeric structure that lacks any additional antenna proteins, like many other cyanobacteria (Hamaguchi *et al*., 2021; Jordan *et al*., 2001; Kato *et al*., 2022; Malavath *et al*., 2018; Xu *et al*., 2021). Therefore, the red-shifted Chls *d* within NIES-2412 PSI are found within the core itself and are analogous to Chl *a* containing PSI red forms.

### Impact of Chl *d* red forms on excitation energy dynamics in NIES-2412 PSI

Two pools of Chl *d* red forms, designated as Red1 and Red2, emitting at ∼745 nm and ∼765 nm, respectively, were identified by the target analysis of the TRF data. The SAS of Red2 is broad, a feature also observed in the 300 ps component from the 77 K TRF global analysis (**Fig. 2D**), and the 750 nm band from the Gaussian deconvolution of the 77 K absorption spectrum (**Fig. S3B**). The large width of these bands is indicative of extensive CT character in the Red2 pool (Novoderezhkin et al., 2016; Romero *et al*., 2009; Sláma *et al*., 2023). Conversely, the SAS of Red1 is relatively narrow, suggesting mainly excitonic character for this red form.

At 77 K, trapping from Red1 and Red2 occurs with distinct lifetimes of ∼50 ps and ∼300 ps, respectively, suggesting that they are not functionally connected at cryogenic temperatures. This implies that the Red1 and Red2 pools are not in direct contact and energy transfer between them at RT occurs via Chls with higher energy. At RT, trapping occurs with lifetimes of ∼16 and ∼60 ps, resulting in an average trapping lifetime of ∼45 ps. The broader nature of the 60 ps DAS compared to the 16 ps DAS, suggests they represent trapping from the Red2 and Red 1 state, respectively, and trapping from these red forms occurs before equilibration is complete.

The target analysis also shows that trapping from the bulk Chls *d* occurs in ∼20 ps, which is very close to a wide variety of Chl a-containing cyanobacterial PSI complexes (Croce and van Amerongen, 2013; Gobets *et al*., 2001) and to the main trapping component that is observed in MBIC11017 PSI (Oliver et al., 2025). The average trapping time of ∼45 ps in NIES-2412 equates to a charge separation efficiency (ϕ_CS_) of ∼98% (where ϕ_CS_ =1 - τ_CS_/τ(2 ns), τ_CS_ is the average trapping lifetime and τ(2 ns) is the excited state lifetime of Chl in the absence of photochemistry (Belgio et al., 2012)). While the incorporation of the Red1 and Red2 pools in NIES-2412 slows down its trapping lifetime, its charge separation efficiency remains high.

### Red form assignment in NIES-2412 PSI

The target analysis of the TRF data, and the Gaussian deconvolution of the NIES-2412 PSI low temperature absorption spectrum show that ∼5 Chls *d* make up the Red 1 pool and 1-2 Chls *d* make up the Red 2 pool. In order to assign these red forms, we resolved the NIES-2412 PSI structure using cryo-EM, and compared this to the structures of MBIC11017 PSI, which does not contain the Red1 and Red2 pools.

The newly resolved B31-B32-B33 trimer and the B20-B41 dimer, both bound by PsaB and interacting with PsaX2 (**Fig. 5**), are strong candidates for these red forms, due to their absence in the MBIC11017 PSI structures (Hamaguchi *et al*., 2021; Xu *et al*., 2021). The NIES-2412 PSI CD spectrum shows a strong negative band at 740 nm (**Fig. 1C**), and the Red1 SAS (emitting at 745 nm, (**Fig. 3B**) is narrow, both of which indicate that Red1 is mainly exitonic in its nature. Calculation of the electronic coupling of the B31-B32-B33 trimer and the B20-B41 dimer shows that they are both strongly coupled, with their oscillator strength almost entirely distributed on the lowest energy exciton state (**Tab. S3-6**).

In the case of the B31-B32-B33 trimer, an almost identical Chl *a* trimer was first identified in the crystal structure of PSI from *T. elongatus (Jordan et al., 2001)* (**Fig 5E,F,G**), which has subsequently been suggested as one of its red forms (Byrdin *et al*., 2002; Toporik et al., 2020). PSI from *Synechocystis* sp. PCC 6803 (*Synechocystis*) possesses only a dimer in this position (Malavath *et al*., 2018), as its PsaB does not possess the loop that binds the third Chl *a*. Examination of the loop residues that co-ordinate the third Chl *d* of the trimer in NIES-2412 PSI reveals an almost identical sequence to the binding loop in *T. elongatus* (**Fig. 5G**), strongly suggesting that the B31-B32-B33 Chl *d* trimer is a red form. Considering that the lowest exciton state of the B31-B32-B33 trimer has an oscillator strength of ∼3, and the Red2 state is formed by 1-2 Chls *d*, we tentatively assign the trimer to the Red1 pool.

In the case of the B20-B41 dimer, comparison with other complexes is not possible, as a Chl at the B41 position is not observed in other cyanobacterial PSI structures. We thus speculate that the B20-B41 Chl *d* dimer is responsible for the Red2 state. However, we acknowledge the possibility that it instead is part of the Red1 pool.

Regardless of the assignment of the B20-B41 dimer to either the Red1 or Red2 pool, there remains 2 additional red Chls *d* to be assigned. Various dimers have previously been considered candidates for the red forms in other cyanobacteria, such as: the A12-A14 dimer (Jiang *et al*., 2025; Kato *et al*., 2022), the B07-A32 dimer (Khmelnitskiy *et al*., 2020), and the B37-B38 dimer (Nurnberg *et al*., 2018; Tros *et al*., 2021). In MBIC11017, Chls *d* at these positions are found within at least one of the two previous structures, preventing us from unequivocally assigning them to red forms in NIES-2412 PSI. Interestingly, we observed sequence differences between MBIC11017 and NIES-2412 in the surrounding regions of these Chl *d* dimers, and the *A. marina* specific PsaI Chl lies very close to the B07-A32 dimer (**Fig. S9A**), but the effect of these changes on the absorption properties is difficult to predict. Indeed, small differences around the Chl 603-609 dimer in the plant light harvesting proteins, Lhca1 and Lhca4, are sufficient to redshift its emission (at 77 K) from 690 nm, in Lhca1, to 732 nm, in Lhca4 (Croce et al., 2002). Moreover, the assignment of red forms in PSI remains challenging, even in well-studied Chl *a* systems such as *Synechocystis* and *T. elongatus*, as they often involve charge-transfer states, requiring detailed spectroscopic and theoretical analyses.

### The role of PsaX2

The presence of a PsaX-like subunit in the NIES-2412 PSI complex is surprising, as it has previously only been found in PSI from thermophilic cyanobacteria (Gisriel et al., 2020; Jordan *et al*., 2001; Koike et al., 1989). Sequence alignment of PsaX suggests a broader yet limited distribution, where it is missing in all basal cyanobacteria except for *Thermosynechococcus* spp. (Jiang *et al*., 2025), which is part of the order Acaryochloridales, and therefore, a somewhat close relative of *Acaryochloris* spp. In MBIC11017, the absence of PsaX in its PSI complex was suggested to increase its flexibility, preventing regions of PsaB from being resolved in its structure (Hamaguchi *et al*., 2021). Indeed, these regions were resolved in NIES-2412 PSI, which may be the reason why an additional 17 – 23 Chls *d* were found compared to MBIC11017 (Hamaguchi *et al*., 2021; Xu *et al*., 2021).

PsaX2 in NIES-2412 is divergent from PsaX found within Chl *a* PSI, but a low level of sequence identity and similarity can be noted in sequence alignments. The NIES-2412 PsaX binds in the same position as that of PsaX in *T. elongatus*, only somewhat tilted relative to each other. However, unlike *T. elongatus*, PsaX2 does not interact with a phospholipid. Instead, PsaX2 provides H-bonds to both the B41 and B32 Chls *d*, both of which are components of the red forms. Therefore, we suggest that PsaX2 not only stabilises the complex, but also modulates the energies and electronic couplings between Chls *d*.

### Comparison with other *A. marina* strains

Investigation into other *A. marina* strains has unveiled significant spectral variations compared to MBIC11017. Many strains show a red-shifted shoulder in the Q_Y_ absorption region (Kiang et al., 2022; Miller et al., 2022; Miller et al., 2005; Mohr et al., 2010), and significantly red-shifted fluorescence emission peaks (Akimoto et al., 2006; Mohr *et al*., 2010; Ulrich *et al*., 2024). Isolated PSI complexes from *A. marina* strains MU05 and P4 (Ulrich *et al*., 2024), and NBRC 102871 (Nagao et al., 2024), display a red-shifted shoulder to their Q_Y_ absorption peak, and red-shifted 77 K emission peaks at ∼750 nm, very similar to NIES-2412 PSI.

Comparison of PsaB from NIES-2412 with other *Acaryochloris* strains reveals that the residues responsible for co-ordinating and interacting with the red forms in NIES-2412 (B20-B41 and B31-B32-B33) are conserved in all compared strains, including the aforementioned *Acaryochloris* sp. NBRC 102871 (Nagao *et al*., 2024). The only exceptions are MBIC11017 (and the closely related strain, *A. marina* MBIC10699), whose PsaB sequences lack many of the residues associated with binding of these red forms (**Fig. S11**). Furthermore, PsaX2, which interacts with both of our proposed red forms in NIES-2412, was not found within the MBIC11017 PSI structures, or its genome, but was present within the genome of all *Acaryochloris* strains that possess the red form binding motifs (and for which a full genome is available) (**Fig. S12**), demonstrating a strong correlation. We therefore hypothesise that the PSI complexes of all *Acaryochloris* strains compared are red-shifted in a similar way to that of NIES-2412, with the exception of MBIC11017 and *A. marina* MBIC10699.

## Methods

### Cell growth

*Acaryochloris marina* MBIC11017 was obtained from the Biological Resource Center, NITE (NBRC; Japan). *Acaryochloris marina* NIES-2412 was obtained from the Microbial Culture Collection at the National Institute for Environmental Studies (MCC-NIES). Cells were grown photoautotrophically in IMK medium with 3.6% (w/v) artificial seawater (Aquaforest), at 25°C and at a constant irradiance of 20 μE m^−2^ s^−1^ using ‘warm white’ LEDs.

### PSI isolation

Thylakoid membranes from MBIC11017 and NIES-2412 were isolated in a similar manner to that previously described (Chen et al., 2005), with some minor changes which are described below. Cells were centrifuged (6500 xg, 15 min) and washed in Buffer A (50 mM MES, 1 M betaine monohydrate, 5 mM CaCl_2_, 5 mM MgCl_2_ and 10% (v/v) glycerol, pH 6.5 adjusted with NaOH). Cells were centrifuged again and resuspended in Buffer A containing EDTA-free protease inhibitor cocktail tablets (cOmplete, Roche), 0.2% (w/v) BSA and 50 μg mL^-1^ DNase I. Cells were lysed by three passages through a pre-chilled French pressure cell at 150 MPa and then centrifuged (2000 xg, 10 min, 4 °C) to remove cell debris. Thylakoid membranes were pelleted by ultracentrifugation (190,000 xg, 30 min, 4 °C) and resuspended in Buffer B (50 mM MES, 1 M betaine monohydrate, 20 mM CaCl_2_, 5 mM MgCl_2_ and 10% (v/v) glycerol, pH 6.5 adjusted with NaOH). Membranes were washed once in Buffer B and resuspended at a Chl *d* concentration of ∼1 mg mL^-1^.

Thylakoids were solubilized for 45 minutes in Buffer B containing 1% (w/v) n-dodecyl-β-maltoside. Unsolubilized material was removed by centrifugation (17,000 xg 10 min at 4 °C). Solubilized thylakoids were loaded on a sucrose density gradient made by freezing and thawing 0.6 M sucrose, 50 mM MES-NaOH at pH 6.5, 10 mM CaCl2, 5 mM MgCl2 and 0.04% (w/v) n-dodecyl-β-maltoside buffer and separated by ultracentrifugation at (270,000 xg, 15 hrs, 4 °C). The PSI trimer band was collected and underwent a second round of sucrose density gradient and ultracentrifugation as specified above (**Fig. S1**).

### Immunoblotting

To verify the purity of NIES-2412 PSI, it was loaded onto a 12% tricine SDS Page Gel (Schagger, 2006), at a concentrations of 0.5 µg of Chl. After electrophoresis, proteins were immunoblotted against PsaB and CP47 antibodies (Agrisera, Sweden) as previously described (Hu et al., 2023). Blots can be seen in **Fig. S1**.

### Steady-state spectroscopy

Absorption spectra were acquired using a Varian Cary 4000 UV-VIS spectrophotometer. Low-temperature absorption spectra at were obtained using a custom-built liquid nitrogen-based cooling system, for which the samples were supplemented to 70% (v/v) glycerol to prevent the formation of ice crystals. For measurements on whole cells, an integrating sphere module was employed. Emission spectra were recorded using a HORIBA JobinYvon-Spex Fluorolog 3.22 spectrofluorimeter, with sample concentrations maintained at an optical density below 0.05 cm^-1^ at the Q_y_ peak to avoid reabsorption. For 77 K emission measurements, samples were frozen in liquid nitrogen and measured in a 1-mm path-length Pasteur pipette. Circular dichroism (CD) spectra were recorded using a Chirascan CD spectrophotometer.

### Stepanov analysis

The fluorescence spectrum of the NIES-2412 PSI complex was calculated from its absorption spectrum using the Stepanov relation (Stepanov, 1957):

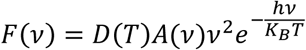

In which F(ν) is the fluorescence spectrum, D(T) is a term that only depends on the temperature (its precise meaning is given in (Stepanov, 1957)), A(ν) is the absorption spectrum, h is the Planck constant, K_B_ is the Boltzmann constant and T is the temperature.

### Gaussian deconvolution

To perform the Gaussian deconvolution of the NIES-PSI 77 K Q_y_ absorption spectrum first the second derivative of this absorption spectrum was computed. The negative peaks of this absorption were then used to guide the peak positions of the individual Gaussians during the decomposition. Specifically, during the fitting, the peak positions of the Gaussians were allowed to deviate two nanometers from the determined second derivative minima, except for the blue region of the spectrum where two Gaussians were needed to accurately fit the data.

### Streak camera measurements

The streak camera setup has been described in detail previously (Hu *et al*., 2023). Excitation pulses were provided at 400 nm at a repetition frequency of 250 kHz at an intensity of 0.8 nJ per pulse. The laser beam was focused within the sample to a spot size of approximately 100 μm. For the RT measurements the sample was constantly stirred using a magnetic stirring bar. For the measurements at cryogenic temperatures the samples were frozen in liquid nitrogen and measured in a Pasteur pipette. For the RT measurements a streak time-window of ∼400 ps was chosen, yielding a time-resolution of ∼7 ps. For the 77 K measurement the sample was measured both using the ∼400 ps time-window and a ∼1.5 ns time-window, and analysed simultaneously in a global analysis (*vide infra*).

### Global and target analysis

Global and target analyses have been performed on the time-resolved fluorescence data using the pyglotaran Python package(van Stokkum et al., 2023; Weißenborn et al., 2023). In the global analysis the time-resolved datasets ψ(λ,t) are fitted with a sum of exponential components that decay separately with a rate k, and which are convolved with the instrument response function (IRF(λ,t)), yielding the decay-associated spectra (DAS(λ)):

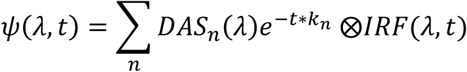

The IRF was modelled as a Gaussian and its FWHM was a free fitting parameter.

The kinetic scheme for the target analysis is presented in Fig. 4A. To estimate the initial excitation input vectors for the separate compartments the areas of the SAS were constrained to be equal(Snellenburg et al., 2013). In addition, the input vectors of the Red1 and Red2 compartment were forced to be minimally 2% of the Bulk input vector. Furthermore, to retrieve kinetics of the pure spectral species, the SAS of the Red1 and Red2 compartment were forced to zero for λ < 720 nm.

From the target analysis results and the total number of Chls per PSI, one can estimate the total number of Chls that are represented by the Bulk, Red1 and Red2 compartment. The ratio of forward/backward rate from the bulk to one of the Red compartments is connected to the Gibbs free energy difference of the two compartments through the Boltzmann equation:

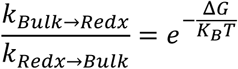

In which the Gibbs free energy is the sum of an enthalpic – and entropic term: ΔG = ΔH -TΔS. The enthalpic energy difference between the compartments is then extracted as the difference in energy that corresponds to the wavelength maxima of the associated SAS, which allows to determine the entropic energy difference between the compartments. The number N in one of the Red compartments is then computed as 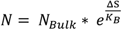.

### Electronic coupling and excitonic dipole distribution calculation

Electronic couplings were calculated using the TrEsp method(Madjet et al., 2006), in combination with the empirical screening expression that was derived from (Chl *a*-containing) Photosystem I trimers by Eder & Renger (2024)(Eder and Renger, 2024) and the transition electrostatic potential charges for Chl *d* as derived by Kimura et al. (2022)(Kimura et al., 2022). To retrieve the exciton levels and connected dipole strength for the triad Chls, we first diagonalized the exciton Hamiltonian, assuming equal site energies for each Chl. This yields the exciton energy levels in the form of the eigenvalues and the wavefunction coefficients C_Jm_ of the individual sites m to each particular exciton J as the eigenvectors. The excitonic transition dipole moments µ_J_ can then be calculated as:

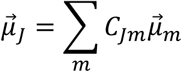

The transition dipole moments of the individual Chls µ_m_ are extracted from the structure as the line connecting the NB and ND nitrogen of each Chl.

### Genome sequencing

To retrieve amino acid sequences for building the CryoEM model, the genome of NIES -2412 was sequenced using short-read next-generation sequencing technologies. Genomic DNA from NIES-2412 was extracted using the Quick-DNA Fungal/Bacterial Miniprep Kit (Zymo Research, Irvine, CA, USA). Whole genome sequencing was performed using the Illumina NovaSeq X plus platform and 150bp pair-end sequencing (Novogene, Cambridge, UK). A total 3.0 GB of data was generated, of which 99.39% were effective reads. Raw filtered reads were assembled into a draft genome using Unicycler v0.5.1 with Spades v4.0.0 and using default parameters (Wick et al., 2017). The total length of the assembly was 8.13 Mbp with an estimated coverage of 395x. The number of contigs ≥200bp was 1148, with the longest segment being 273,609 bp and N50 72,230. Genome completeness and contamination was estimated with CheckM v1.2.4 (Parks et al., 2015). Overall, the NIES-2412 genome was found to be 8.05 Mbp with a completeness of 99.0% and 1.7% contamination.

Annotation of the genome was carried out using Prokka v1.14.5 using default parameters (Seemann, 2014) on the KBase webserver (Arkin et al., 2018).

### Grid preparation

UltrAuFoil 2 μm hole size and 1 μm hole spacing with a 300 mesh grid were glow-discharged for 100 s at 25 mA. At a temperature of 4 °C and 100% humidity and in the presence of only a dim green light, 3.5 ul of PSI sample (∼1 mg/ml of Chl concentration) were applied and grids were blotted and immediately plunge-frozen in liquid ethane with a Vitrobot Mark IV (Thermo Fisher Scientific).

### Data acquisition and processing

Micrographs were acquired using a Krios I (Thermo Fisher Scientific) operated at 300 kV and a magnification of 81000x. Images were recorded on a K3 (Gatan Inc.) with a pixel size of 1.058 Å and a dose of 40 electrons per Å^2^. Images were collected in super-resolution mode with a SelectrisX energy filter with a slit width of 20 eV. The targeted defocus range was varied from -0.8 to -2 µm using the EPU software (Thermo Fisher). A total of 12085 movies were collected from a single grid. The frames were aligned, dose weighted, and the contrast transfer function (CTF) was estimated in CryoSPARC v4.5.1 (Punjani et al., 2017). Micrographs were curated by removing those with CTF fits worse than 10 Å and other unsuitable parameters. The subset obtained 77% of the initial micrographs and was used to hand pick ∼500 PSII-type particles across the defocus range. Particles used to train topaz (Bepler et al., 2019) template pick across the entire dataset, yielding 334934 picks. After multiple rounds of topaz training, 2D classification and *ab initio* refinement, one major class emerged, one corresponding to PSI trimers. Duplicated particles were removed from the stack and of ∼20,000 particles was used for homogeneous refinement to obtain an initial map at a resolution of ∼3 Å and to confirm C3 symmetry. After multiple rounds of per particle CTF refinement and local motion correction, a set complete set of 151833 particles was used to perform non-uniform refinement imposing C3 symmetry, producing a map at a global resolution of 2.63 Å based on GS-Fourier Shell Correlation at a cut-off of 0.143. The Guinier’s analysis reports an estimated B-factor of 68.4. To obtain further definition of the PsaX2 subunit, local refinement was performed on symmetry expanded particles. A mask encompassing the other two subunit was used to subtract the signal from the other two monomers. The reconstruction of the PSI monomer has a GS-FSC of 2.44 Å.

### Model building

The trimeric PSI from MBIC11017 (PDB ID: 7COY)(Hamaguchi *et al*., 2021) was fitted to the ESP map using the Phenix software suite(Afonine et al., 2012), mutated to the correct sequence with Chainsaw (Phenix) and refined in Coot(Emsley and Cowtan, 2004). Each subunits density was evaluated amino acid by amino acid and the model was then refined in real space in Phenix.

### Sequence Alignments

Both protein sequences and nucleotide sequences from the NIES-2412 genome were used to search for PSI sequences from the different *Acaryochloris* strains used in this study. With the exception of PsaI and PsaX, protein sequences in the nr database were searched for using blastp(Camacho et al., 2009). In the case of PsaI, sequences were found via tblastn in both the nt and wgs databases, and subsequently translated. In the case of PsaX, sequences were found using a nucleotide input via blastn in both the nt and wgs databases, and subsequently translated. Sequences were aligned using the Muscle algorithm(Edgar, 2004) within SeaView (Gouy et al., 2010) using the default settings. Alignments were visualised using TEXshade(Beitz, 2000).

## Acknowledgements

This work is supported by the Dutch Organization for Scientific Research (NWO) via a TOP grant (714.018.001) and the ERC Advanced grant FARED WELL (101141764) (R.C, E.E, T.J.O). The financial support of a UKRI Future Leaders Fellowship is gratefully acknowledged (MR/T017546/1, MR/T017546/2, and MR/Y011635/1, T.C. and V.C.P). This work is also supported by a Marie Skłodowska-Curie grant agreement (955520 – G.C, A.W.R, A.F), the Biotechnology and Biological Sciences Research Council (BB/R001383/1 – A.W.R, A.F; BB/V002015/1 – A.W.R, A.F; BB/R00921X – A.W.R, A.F; BB/Z516740/1 – G.C, A.W.R, A.F), a Leverhulme Trust Grant (RPG-2022-203 – A.W.R, A.F), a Royal Society Research Professorship (A.W.R), and an Imperial College President’s Scholarship (H.F.L).

Computing resources were provided by the Apocrita High Performance Computing Facility at Queen Mary University of London (http://doi.org/10.5281/zenodo.438045). We thank the Centre for Structural Biology at Imperial College for the training provided, for help with the early stages of sample screening and with data collection. We thank Diamond Light Source for the access and support of the cryo-EM facilities at the UK national electron Bio-Imaging Center (eBIC), proposal BI25127. This project made use of the Computing Platform for Electron Microscopy at Imperial College funded by the BBSRC Mid-range equipment Initiative 22ALERT (BB/X019284/1).

## Author contributions

Conceptualization, T.J.O; Methodology, T.J.O, E.E, G.C, H.F.L, V.C.P, T.C, R.C; Investigation, T.J.O, E.E., G.C, H.F.L; Formal Analysis, T.J.O, E.E, G.C, H.F.L; Writing – Original Draft, T.J.O, E.E, R.C; Writing – Review & Editing, T.J.O, E.E, G.C, H.F.L, V.C.P, A.F, T.C, A.W.R, R.C; Supervision, A.F, T.C, A.W.R, R.C; Funding acquisition, A.F, T.C, A.W.R, R.C

## Data availability

The data that support the findings of this study are available in the Supporting Information of this article. Additionally the source data for **Figs. 1 & 2** can be found at http://doi.org/10.5281/zenodo.16635114. The Whole Genome Shotgun project has been deposited at DDBJ/ENA/GenBank under the accession JBPVIU000000000. The version described in this paper is version JBPVIU010000000 (NCBI Bioproject PRJNA1295459).

