## Supplementary Materials for "Far-red light absorption strategies and their structural basis in Photosystem I of Acaryochloris marina NIES-2412"

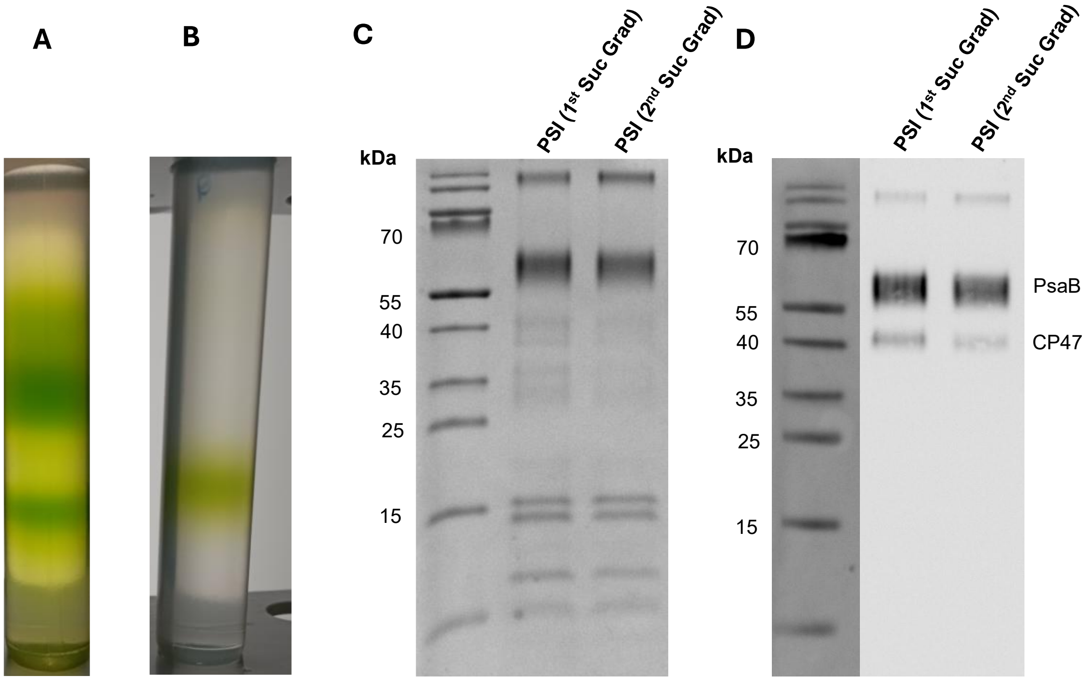


**Figure S1. Purification of *Acaryochloris marina* NIES-2412 Photosystem I. (A)** Sucrose density gradient of NIES-2412 solubilised thylakoids. The lowest major band contained NIES-2412 PSI. **(B)** Sucrose density gradient of cleaned NIES-2412 PSI. **(C)** 12% tricine SDS PAGE gel of NIES-2412 PSI (Lane 1), and cleaned NIES-2412 PSI. Samples were loaded at a Chl concentration of 0.5 µg **(D)** Immunoblot using PsaB and CP47 antibodies to show minimal contamination of PSII in the NIES-2412 PSI. Samples were loaded at a Chl concentration of 0.5 µg.

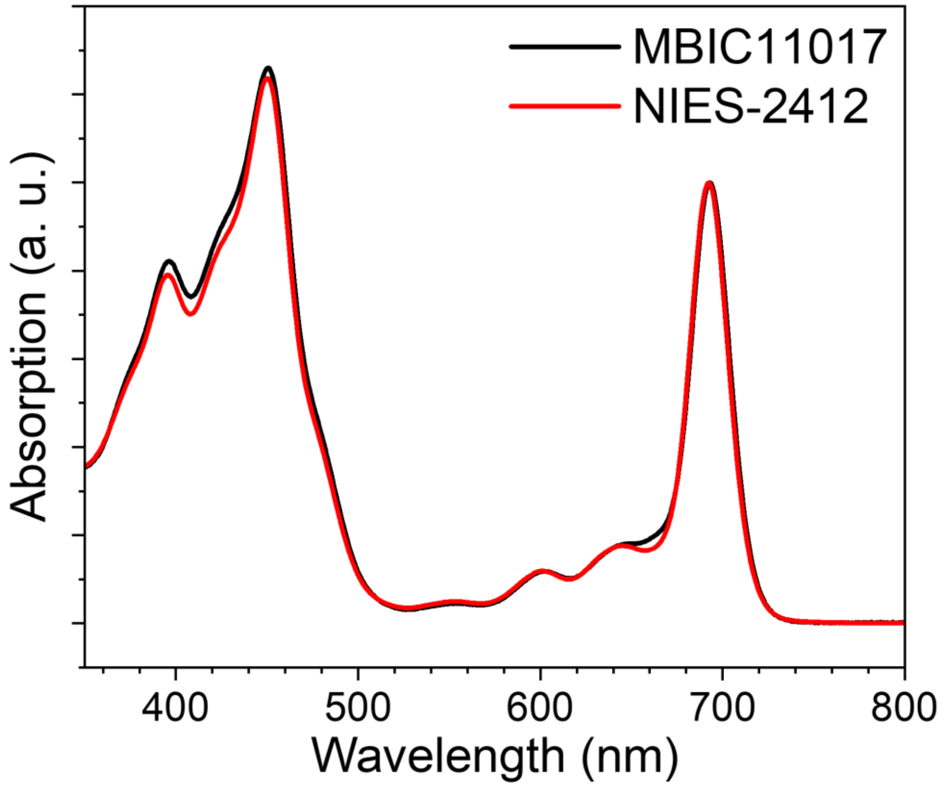


**Figure S2. Absorption spectrum of pigments extracted from Acaryochloris marina NIES-2412 and MBIC11017 PSI.**

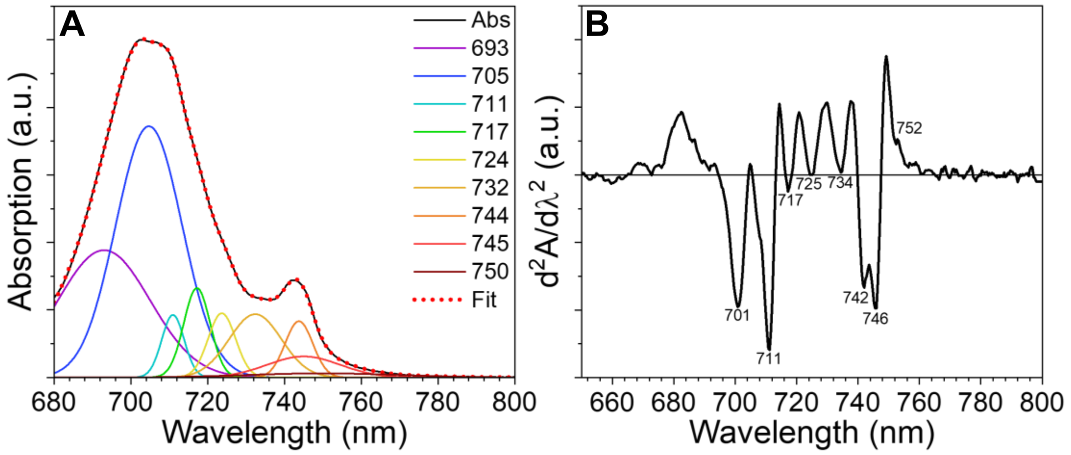


**Figure S3. Gaussian deconvolution and Second-derivative of the NIES-2412 PSI 77 K absorption spectrum. (A)** The Gaussian deconvolution of the NIES-2412 PSI 77 K absorption spectrum was guided by the negative peaks of second derivative spectrum **(B),** in particular: the peak positions of the Gaussians were allowed to deviate two nanometers from the determined second derivative minima, except for the blue region of the spectrum where two Gaussians were needed to accurately fit the data.

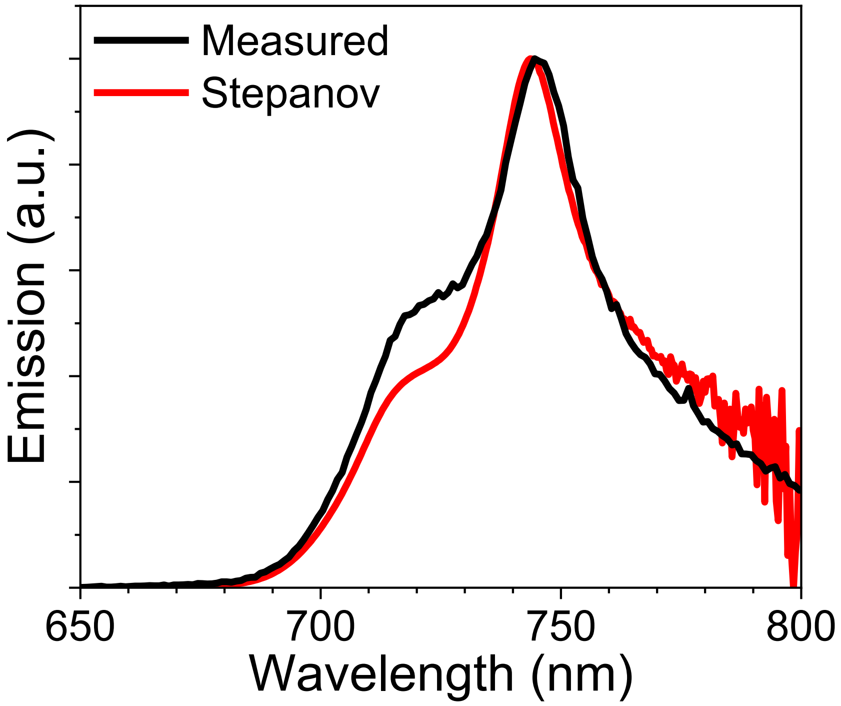


**Figure S4. Measured – and Stepanov predicted RT emission spectrum for the NIES-2412 PSI complex.**

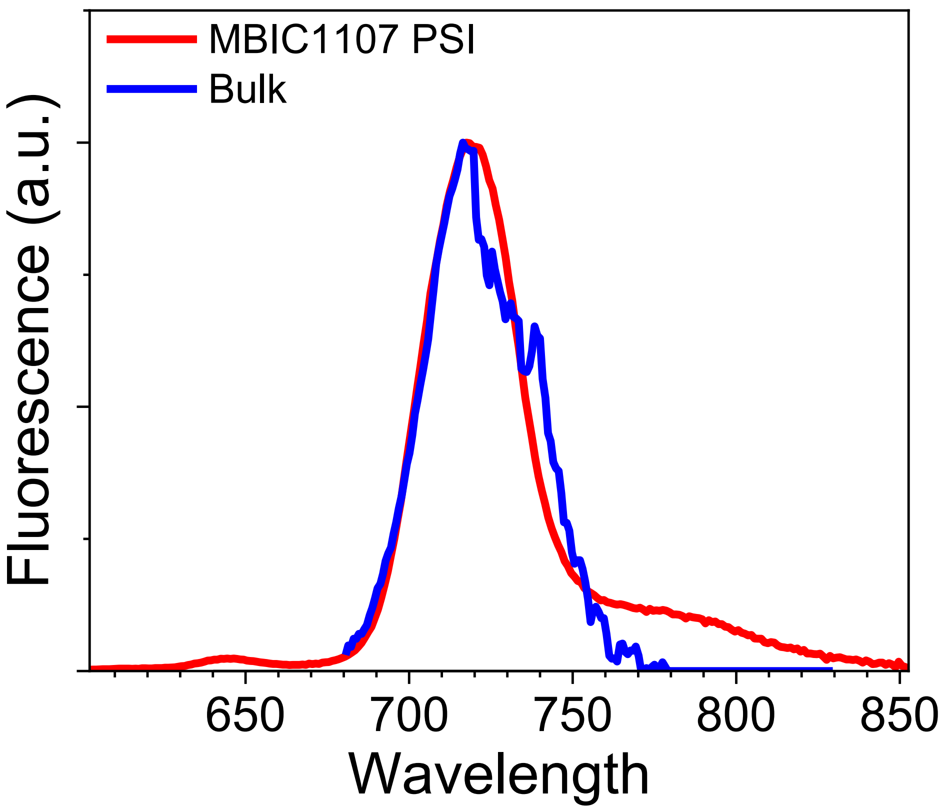


**Figure S5. Comparison of the Bulk compartment SAS from the NIES-2412 PSI target analysis and the emission spectrum of MBIC1107 PSI (Oliver *et al.*, 2025).**

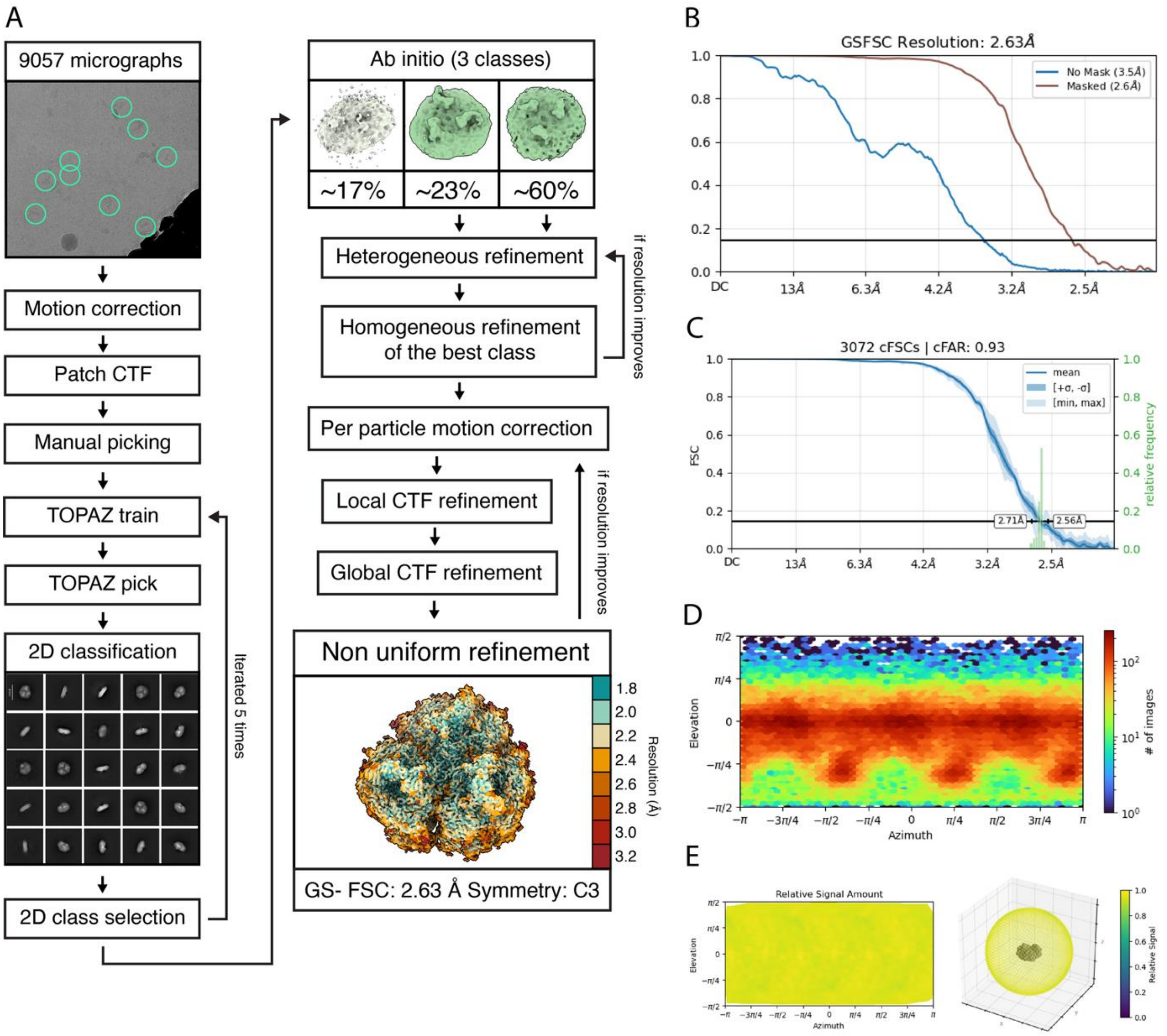


**Fig S6. Processing workflow and reconstruction diagnostic plots for trimeric PSI. (A)** Processing workflow to obtain the final reconstruction of the trimeric PSI. **(B)** GS-FSC curve for the trimeric PSI. **(C)** cFSC curves for the trimeric PSI complex. **(D)** Viewing direction distribution of the particle stack. **(E)** relative signal amount for each of the viewing directions.

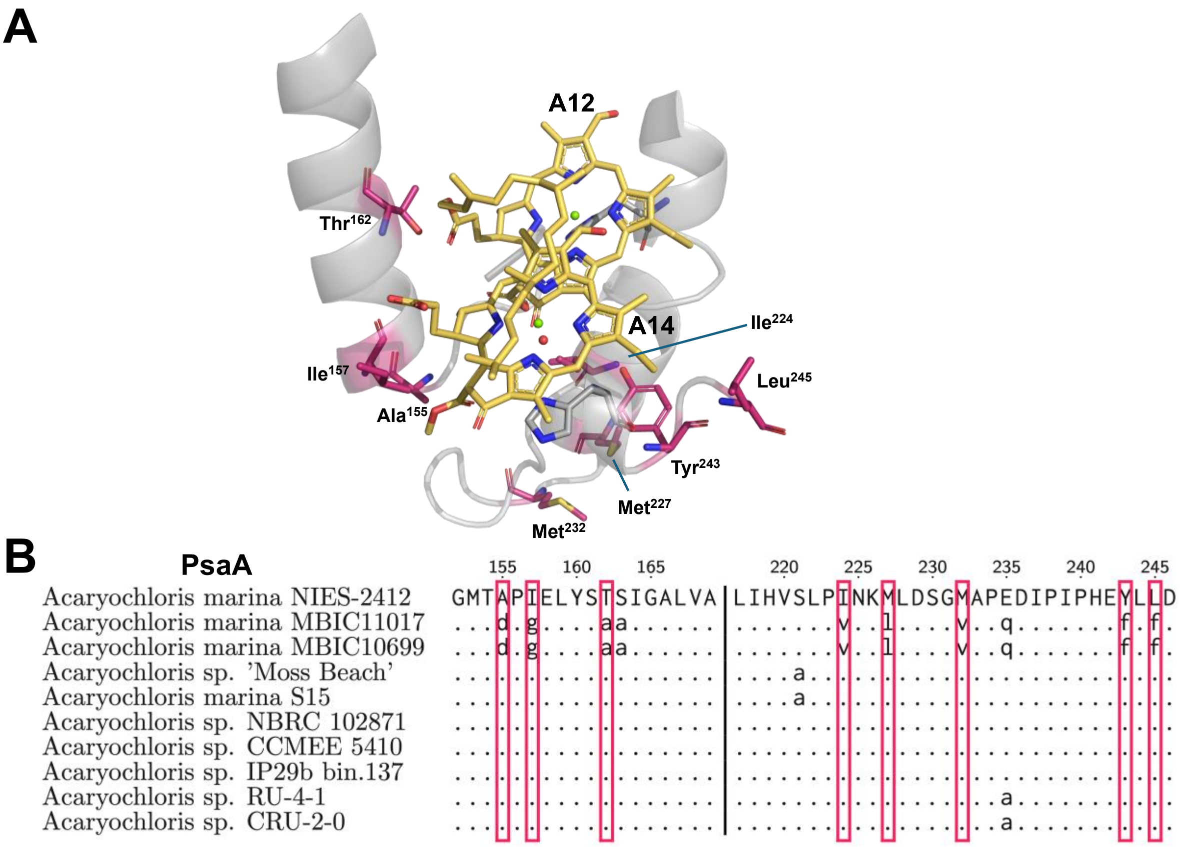


**Figure S7. PsaA differences between NIES-2412 and MBIC11017 PSI around the A12-A14 Chl *d* dimer.** **(A)** The A12-A14 Chl *d* dimer in NIES-2412 PSI. PsaA residues highlighted in pink are different from those found in MBIC11017 PsaA. **(B)** Sequence alignment of PsaA from NIES-2412, MBIC11017 and various Acaryochloris strains. Red boxes highlight the PsaA residues that are different between NIES-2412 and MBIC11017 in the vicinity of the A12-A14 Chl *d* dimer. Dots indicate identical residues to the first sequence in the alignment. Lowercase letters indicate residues that are different to the first sequence in the alignment.

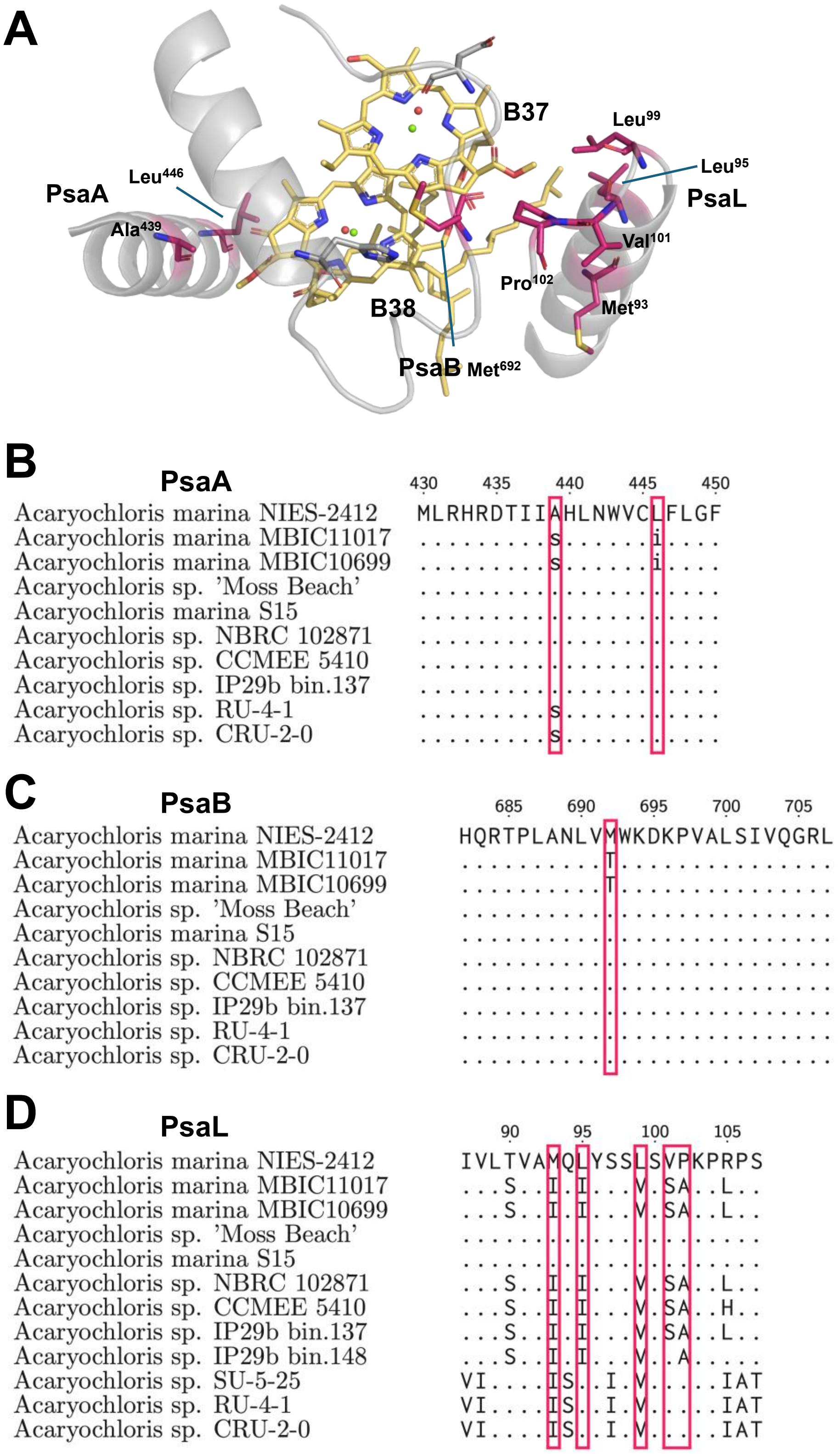


**Figure S8. Differences in PsaA, PsaB, and PsaL between NIES-2412 and MBIC11017 PSI around the B37-B38 Chl *d* dimer. (A)** The B37-B38 Chl *d* dimer in NIES-2412 PSI. Residues highlighted in pink are different from those found in MBIC11017 PSI. Sequence alignments of PsaA **(B)**, PsaB **(C)**, and PsaL **(D)** from NIES-2412, MBIC11017 and various Acaryochloris strains are shown below. Red boxes highlight the residues that are different between NIES-2412 and MBIC11017 in the vicinity of the B37-B38 Chl *d* dimer. Dots indicate identical residues to the first sequence in the alignment. Lowercase letters indicate residues that are different to the first sequence in the alignment.

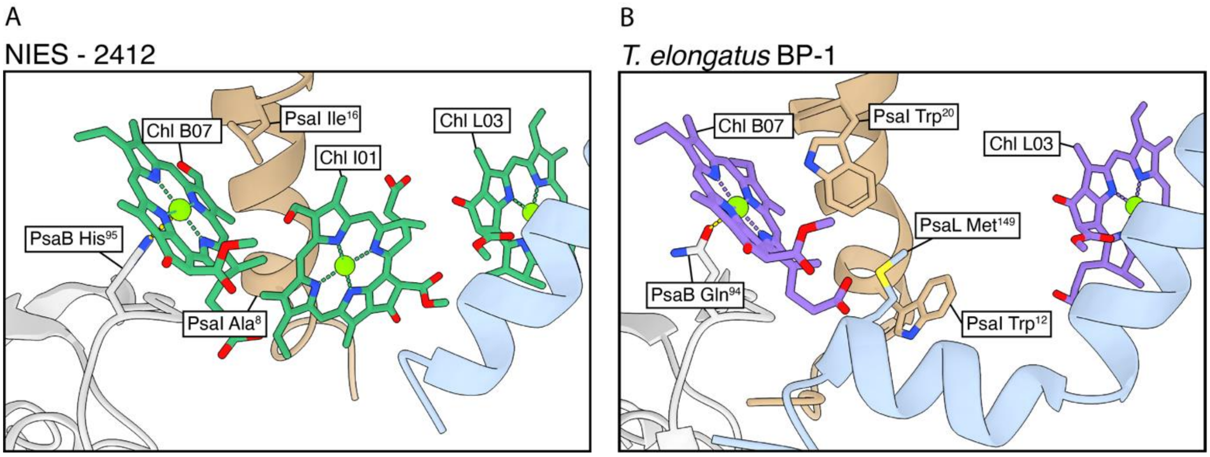


**Figure S9. The location of the additional Chl *d* bound by PsaI Photosystem I (PSI) from *Acaryochloris marina* NIES-2412. (A)** Chl *d* I01 is found between the N-terminus of PsaI and the C-terminus of PsaL in *A. marina* NIES-2412 PSI. **(B)** The equivalent location in (PSI) from *T. elongatus*, demonstrating the lack of a Chl at this position.

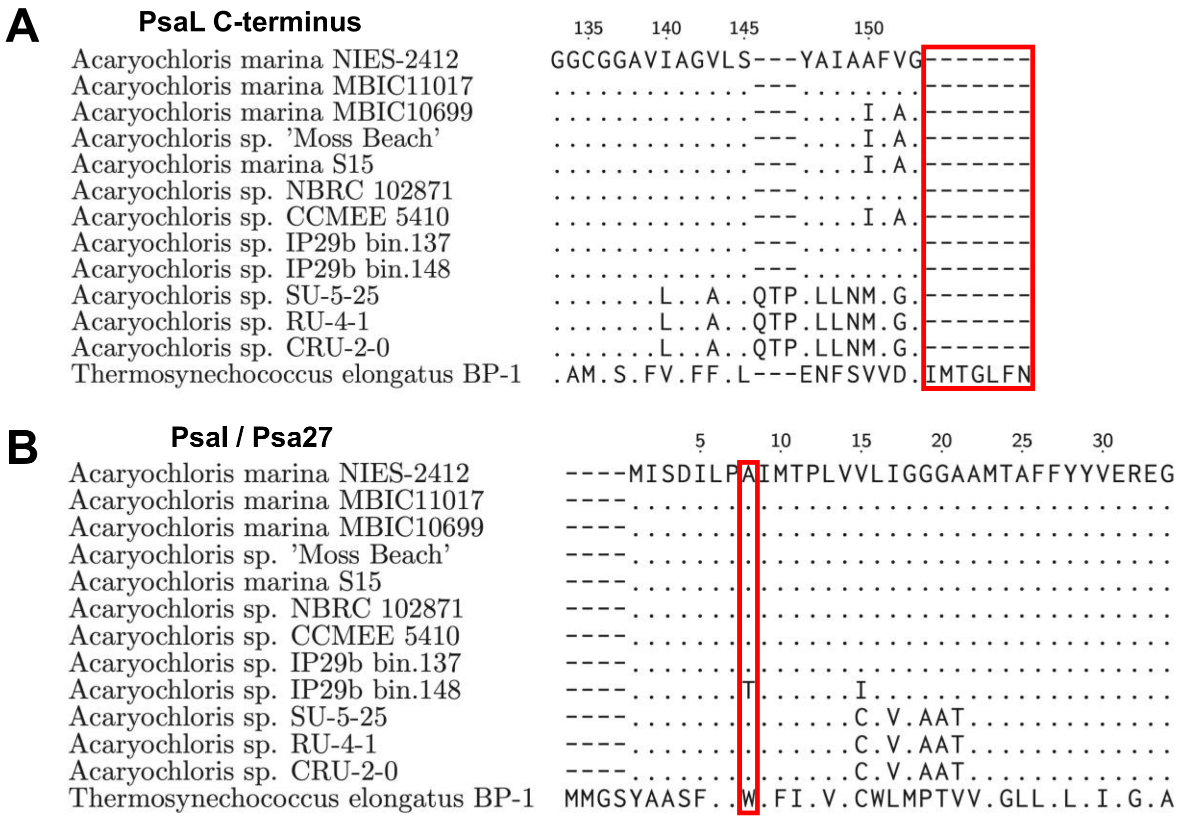


**Figure S10. Sequence alignment of PsaL and PsaI (Psa27) from various *Acaryochloris* strains compared to *Thermosynechococc*us elongatus BP-1. (A)** PsaL C-terminus alignment of NIES-2412, MBIC11017 and various other *Acaryochloris* strains, compared to *T. elongatus*. The red box highlights the truncated C-terminus of PsaL in *Acaryochloris* strains vs. *T. elongatus*. **(B)** PsaI (referred to as Psa27 by Hamaguchi *et al.* (2021) in MBIC11017) alignment of NIES-2412, MBIC11017 and various other Acaryochloris strains, compared to *T. elongatus*. The red-box highlights Ala$^{8}$ in *Acaryochloris* strains which is found as Trp$^{12}$ in *T. elongatus*. Dots indicate identical residues to the first sequence in the alignment. Lowercase letters indicate residues that are different to the first sequence in the alignment.

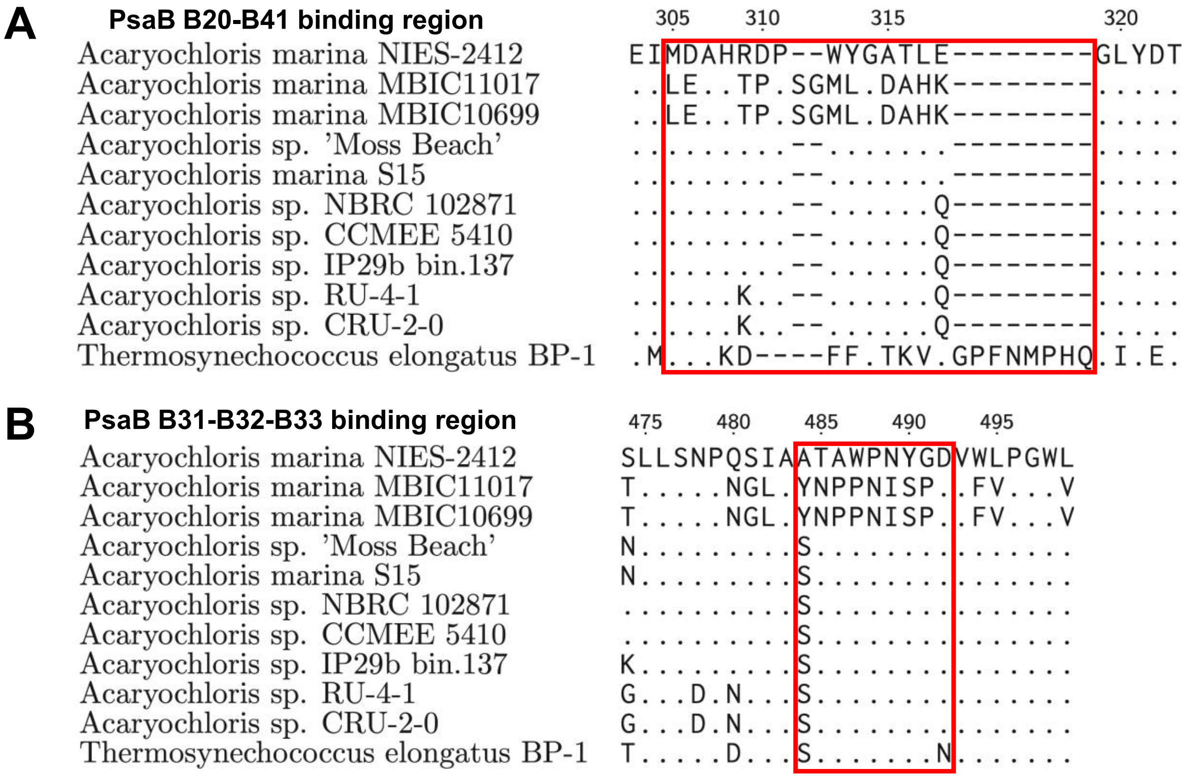


**Figure S11. Sequence alignment of PsaB from various *Acaryochloris* strains compared to *Thermosynechococc*us elongatus BP-1 in the B20-B41 and B31-B32-B33 Chl binding regions. (A)** B20-B41 binding region alignment of NIES-2412, MBIC11017 and various other *Acaryochloris* strains, compared to *T. elongatus*. **(B)** B31-B32-B33 binding region alignment of NIES-2412, MBIC11017 and various other *Acaryochloris* strains, compared to *T. elongatus*. Red boxes highlight the loop regions responsible for binding the additional Chls *d* i.e. B41 and B33, respectively. Dots indicate identical residues to the first sequence in the alignment. Lowercase letters indicate residues that are different to the first sequence in the alignment.

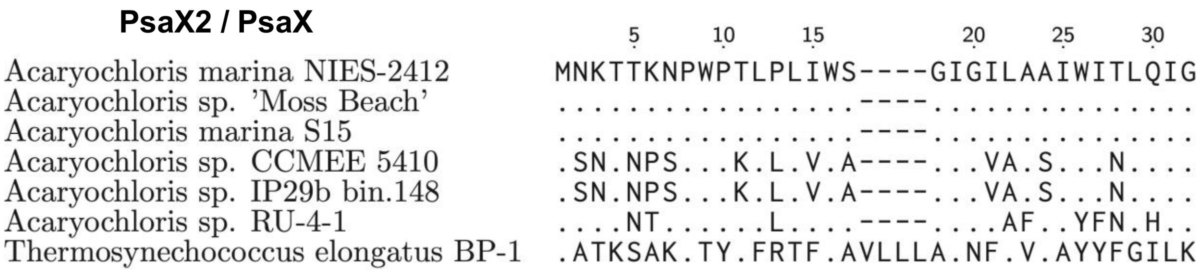


**Figure S12. Sequence alignment of PsaX2 from various *Acaryochloris* strains compared to PsaX from *Thermosynechococc*us elongatus BP-1.** Dots indicate identical residues to the first sequence in the alignment. Lowercase letters indicate residues that are different to the first sequence in the alignment.

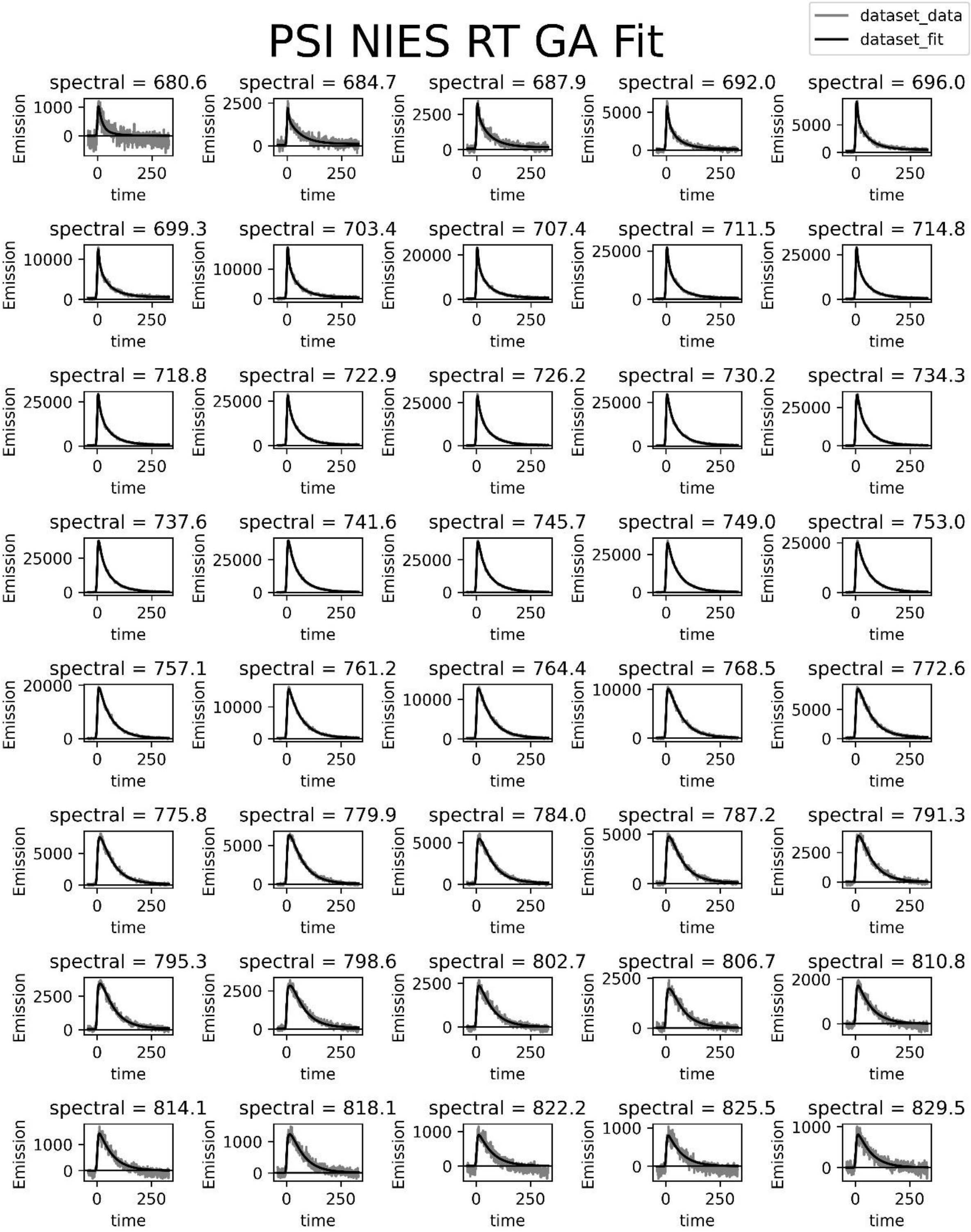


**Figure S13. Fitting results for the global analysis of the RT time-resolved fluorescence experiment on NIES-2412 PSI.**

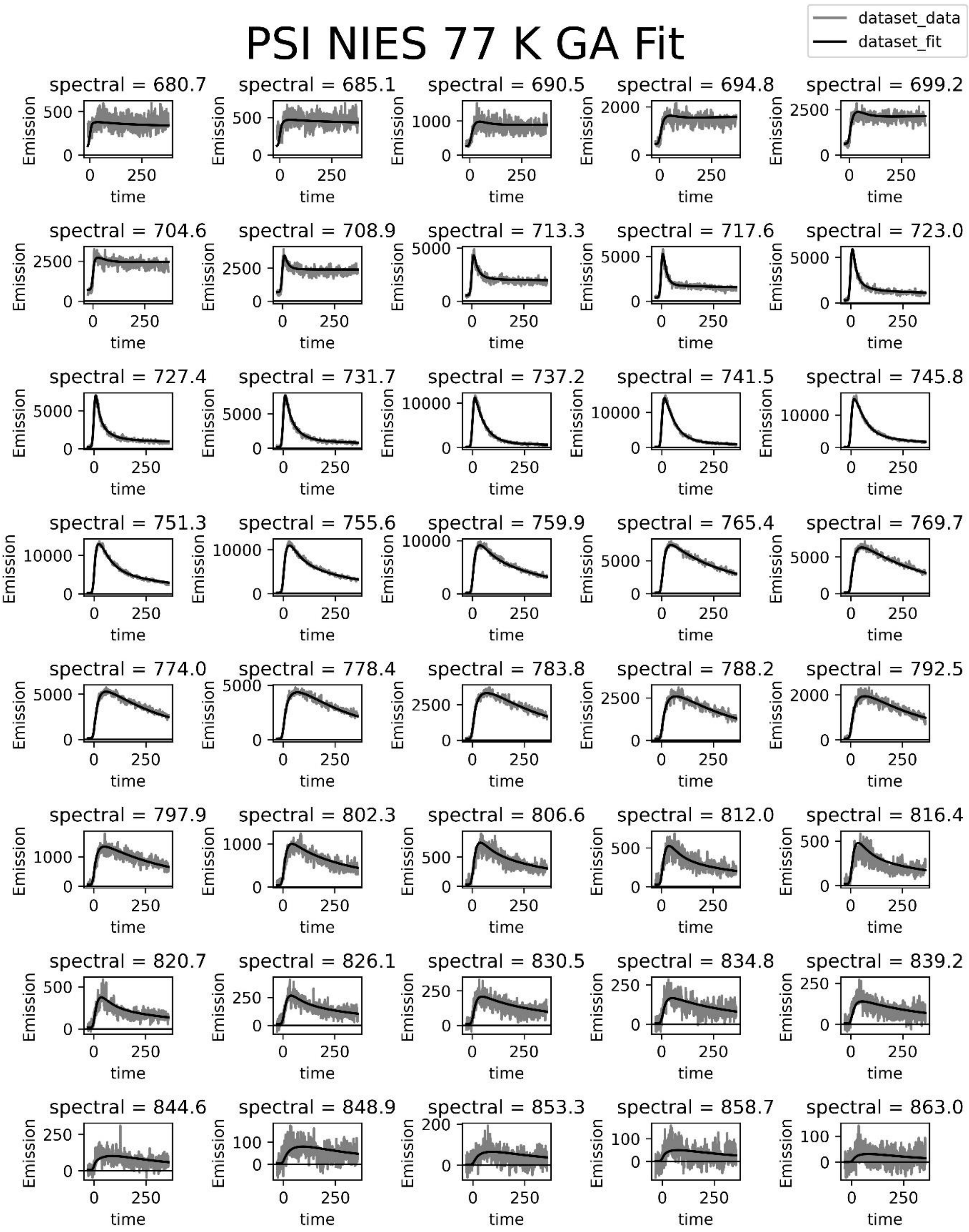


**Figure S14. Fitting results for the global analysis of the 77 K time-resolved fluorescence experiment on NIES-2412 PSI for the ~400 ps time-window data.**

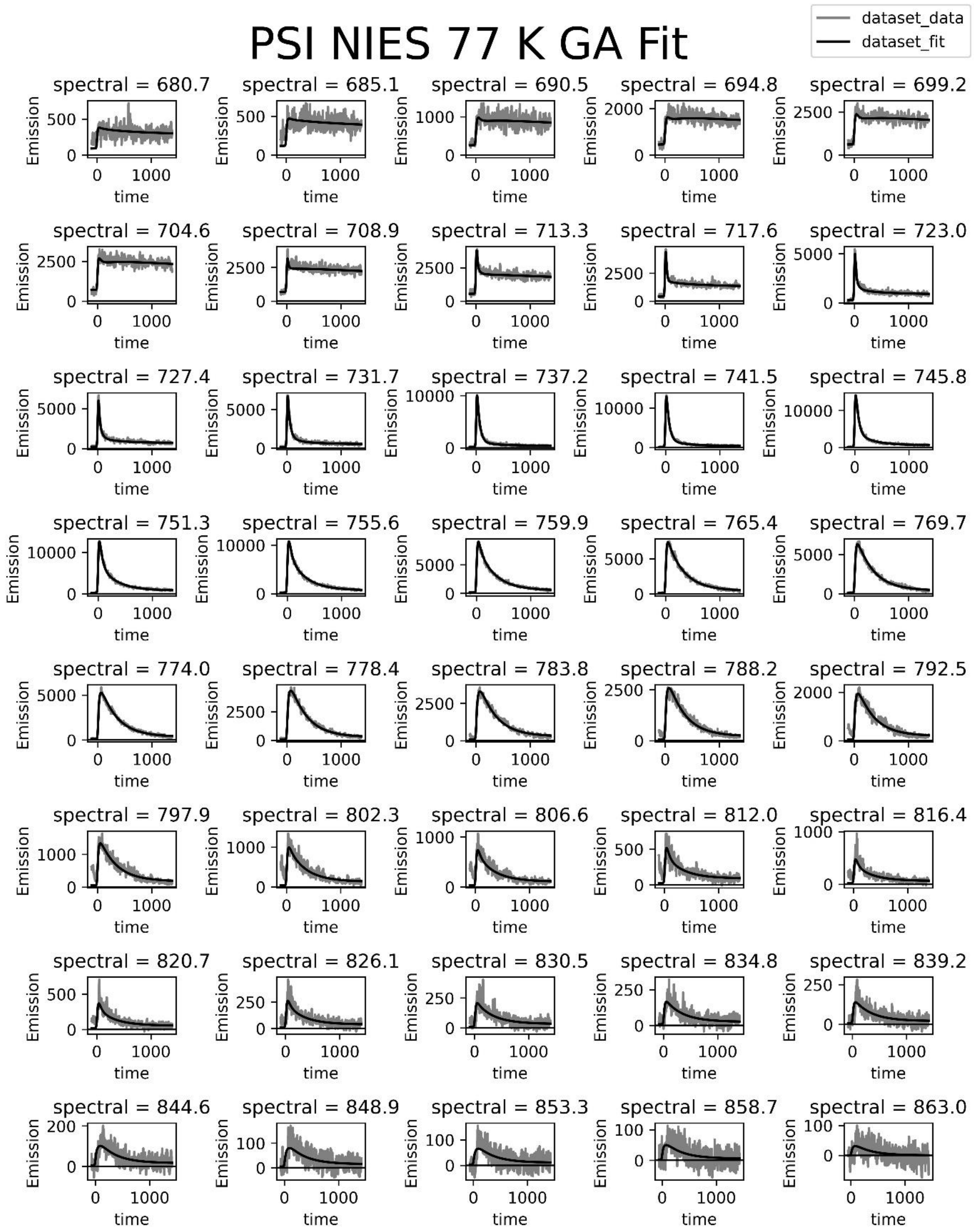


**Figure S15. Fitting results for the global analysis of the 77 K time-resolved fluorescence experiment on NIES-2412 PSI for the ~1.5 ns time-window data.**

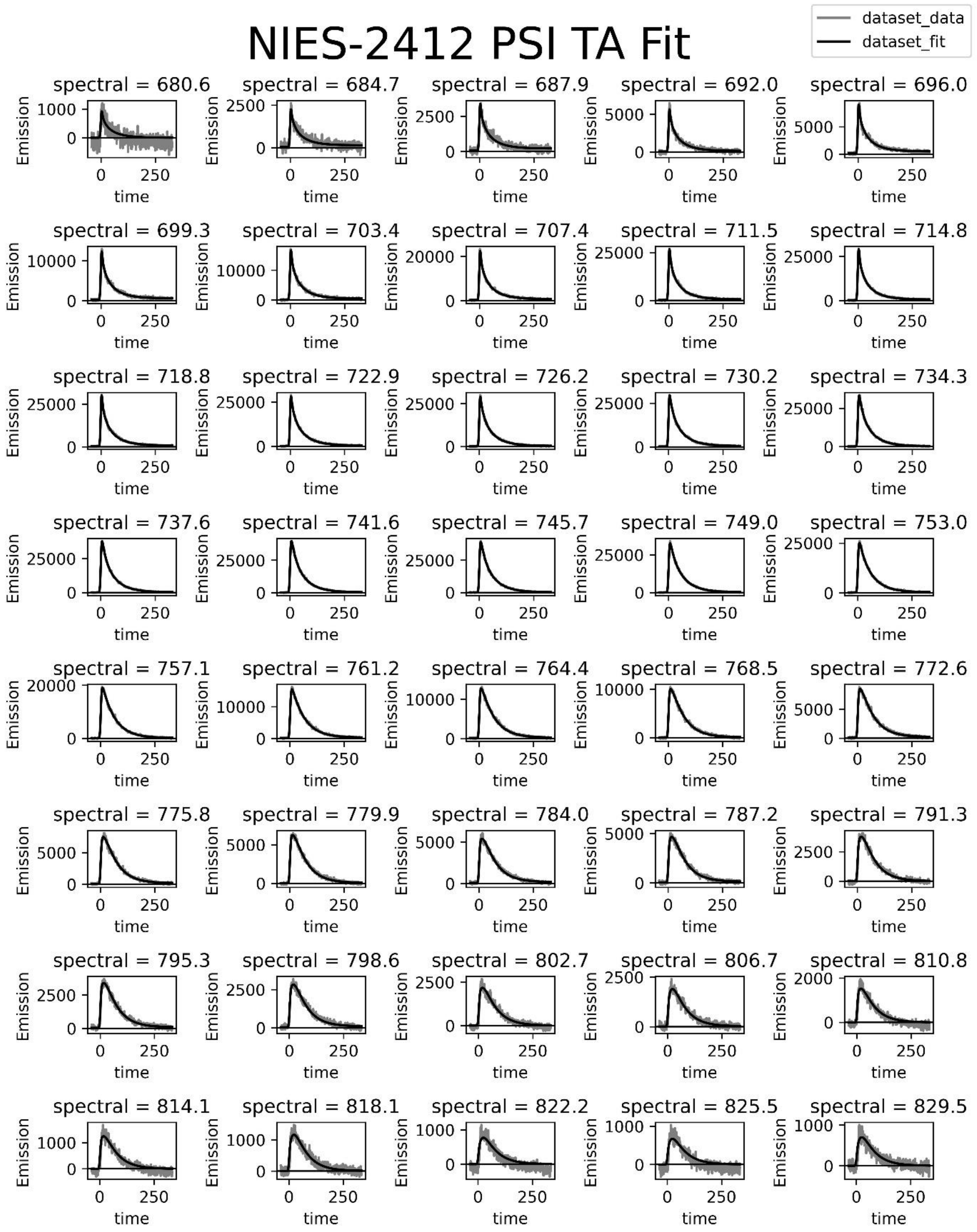


**Figure S16. Fitting results for the target analysis of the RT time-resolved fluorescence experiment on NIES-2412 PSI.**

| | **Bulk** | **Red1** | **Red2** |
|---|---|---|---|
| **$\lambda_{max}$ (nm)** | 716.4 | 744.9 | 757.1 |
| **ΔH ($k_BT$)** | 0 | -2.62 | -3.68 |
| **ΔG ($k_BT$)** | 0 | 0.25 | 0.74 |
| **TΔS ($k_BT$)** | 0 | -2.87 | -4.42 |
| **$N_{Chl}$** | 90 | 5 | 1 |

**Tab S1. Thermodynamic properties of NIES-2412 PSI arising from the target analysis.** The wavelength maxima are retrieved as the maxima of the corresponding SAS of the compartments. These maxima are used to calculate the enthalpic energy difference (ΔH) between the compartments, with the Bulk as a reference. The Gibbs free energy difference (ΔG) between the compartments is calculated from the rate equilibria that arose from the target analysis, and the entropic energy difference (TΔS) follows from the equation ΔH- ΔG = TΔS. The number of Chls in the Red compartments is then calculated using $N_{Chl} = N_{Bulk}\, e^{\frac{\Delta S}{K_B}}$. We have used a temperature of 293 K for the calculations.

### Data collection

| | |
|---|---|
| Microscope | Krios I |
| Camera | K2 |
| Magnification | 81000x |
| Voltage (kV) | 300 |
| Electron exposure (e-/Å$^2$) | 40 |
| Defocus range (μm) | -0.8 to -2.0 |
| Pixel size (Å) | 1.058 |
| Energy filter | Selectris (20 eV) |
| Exposures | 12085 |
| Image format | EER |

### Data processing

| | |
|---|---|
| Box size | 450 px |
| Initial particles (no.) | 334934 |
| Final particles (no.) | 151833 |
| Symmetry | C3 |
| Map resolution (Å) | 2.63 |
| Map sharpening *B* factor | -68.4 |

### Model Refinement

| | |
|---|---|
| Refinement package | PHENIX |
| Initial model used | 7COY |
| Real/reciprocal space | Real Space |
| Resolution cutoff | 2.70 |

### Model Validation

| | |
|---|---|
| MolProbity score | 1.29 |
| ClashScore | 5.35 |
| Bond length R.M.S.D. (Å) | 0.002 |
| Bond angles (°) | 0.555 |
| Poor rotamers | 0.24% |

| | |
|---|---|
| Favored rotamers | 93.48% |
| Ramachandran outliers | 0.05% |
| Ramachandran favored | 98.67% |

**Tab S2. Data collection, processing and model building and validation parameters.**

| | B31 | B32 | B33 |
|---|---|---|---|
| **B31** | 0 | -112 | -25 |
| **B32** | -112 | 0 | -80 |
| **B33** | -25 | -80 | 0 |

**Tab S3. Exciton Hamiltonian for the B31, B32 & B33 Chl triad.** Electronic coupling values are listed in the off-diagonal and the relative site-energies are arranged on the diagonal. All values in $cm^{-1}$.

| Exciton energy level ($cm^{-1}$) | Dipole strength distribution (%) |
|---|---|
| -151 | 95 |
| 24 | 4 |
| 127 | 1 |

**Tab S4. Exciton levels and excitonic dipole strength distributions for the exciton Hamiltonian of Tab. S3.**

| | B20 | B41 |
|---|---|---|
| **B20** | 0 | 122 |
| **B41** | 122 | 0 |

**Tab S5. Exciton Hamiltonian for the B20 & B41 dimer.** Electronic coupling values are listed in the off-diagonal and the relative site-energies are arranged on the diagonal. All values in $cm^{-1}$.

| Exciton energy level ($cm^{-1}$) | Dipole strength distribution (%) |
|---|---|
| -122 | 93 |
| 122 | 7 |

**Tab S6. Exciton levels and excitonic dipole strength distributions for the exciton Hamiltonian of Tab. S5.**
